# Polarization Increases Nuclear Stiffness in Macrophages Despite Reduction in Lamin A/C Levels

**DOI:** 10.1101/2025.11.22.689940

**Authors:** Margaret A. Elpers, Jacob Odell, Sarah J. Henretta, Tong Shu, Yogeshwari Sanjayrao Ambekar, Hassan Saadi, Graeme F. Woodworth, Warren R. Zipfel, Giuliano Scarcelli, Liam J. Holt, Jan Lammerding

## Abstract

Macrophages are innate immune cells contributing to tissue homeostasis and various pathologies. Signals from their environment can lead macrophages to adapt distinct functional phenotypes, a process called polarization. Because macrophages have been previously shown to degrade the nuclear envelope proteins lamin A/C upon pro-inflammatory polarization, and lamins are considered key determinants of nuclear deformability, we aimed to address the effect of pro-inflammatory stimulation on nuclear mechanics. We present the surprising finding that polarized bone marrow-derived macrophages have less deformable nuclei than unpolarized macrophages, despite their reduced lamin A/C levels. Furthermore, pro-inflammatory macrophages exhibited altered chromatin dynamics relative to unpolarized macrophages, including redistribution of trimethylated histone H3K9 (H3K9me3) from the nuclear periphery to the interior and increased chromatin compaction. Our findings suggest a model in which pro-inflammatory stimulation of macrophages induces chromatin changes that drive nuclear stiffening, and that in these cells, chromatin, rather than the nuclear lamina, is the major driver for resisting nuclear deformation. These findings may have functional relevance for the physiological function of polarized macrophages, as the mechanical properties of the nucleus can influence how these cells adapt and respond to their environments in the context of cell migration or inflammatory disease pathologies.

## Introduction

The mechanical properties of the nucleus determine its resistance to deformation in response to cytoskeletal and extracellular forces, such as actomyosin mediated intracellular tension^1–3^, tissue contraction or stretching^4,5^, and during cell migration through confined spaces^6–8^. For example, nuclear deformability governs the ability of cells to successfully migrate through interstitial spaces on the scale of 1-3 µm in diameter, which is essential in developmental processes^9^, cancer metastasis^10,11^, inflammation, and wound healing^6,12,13^. Furthermore, since nuclear deformation can induce various cellular signaling pathways, nuclear envelope rupture, DNA damage, and changes in gene expression and chromatin organization^14^, the mechanical properties of the nucleus contribute to numerous physiological functions such as differentiation^15^ and inflammation^16^, and are also implicated in disease pathologies including cancer metastasis^10^ and muscle related diseases such as laminopathies^17,18^. Additionally, nuclear deformability affects how cells sense and respond to mechanical cues and physical confinement through a process known as mechanotransduction, in which the nucleus activates downstream pathways in response to alterations in increased nuclear membrane tension, allowing cells to adapt their functions such as cell contractility and migration to their physical microenvironment^19–21^.

In many cell types, including fibroblasts, muscle cells, and cancer cells, the resistance of the nucleus to large deformations is primarily determined by the nuclear lamina^14,22,23^, a dense filamentous protein meshwork underlying the inner nuclear membrane primarily comprised of lamins^24^. Lamins are intermediate filament proteins that provide structure and support to the cell nucleus, with both A-type (lamins A/C, encoded by the *LMNA* gene) and B-type (lamins B1/B2, encoded by *LMNB1* and *LMNB2*, respectively) lamins contributing to nuclear morphology^25,26^, chromatin organization^27–29^, and gene expression^30^. Cells with lower lamin levels, particularly with reduced lamin A/C expression, have substantially more deformable nuclei^8,15,22,31,32^, and lamin A/C levels of different cell types positively correlate with the stiffness of the corresponding tissue^33^. Besides lamins, other nuclear envelope proteins, such as emerin^34,35^ and LAP2β^36^, and chromatin in the nuclear interior contribute to nuclear mechanical stability and resistance to deformation.

Experiments in which nuclei isolated from mouse embryo fibroblasts were stretched at physiological strain rates found that chromatin was particularly crucial to resist small nuclear deformation,^22^ and that chromatin histone modifications regulate nuclear stiffness independent of lamins^37^. It is now well recognized that changes in chromatin organization can substantially alter nuclear mechanics. For example, altering chromatin compaction using osmotic swelling or shrinking, upregulation of nucleosome binding proteins, or treatment with histone deacetylase inhibitors modulates nuclear deformability^15,22,23,38^, and histone modifications and the abundance of heterochromatin and euchromatin contribute to the mechanical properties of the nucleus15,37,39,40.

Although the physical properties of the nucleus have been extensively characterized in numerous cell types, the nuclear mechanics of immune cells remain understudied and underappreciated. Here, we focus on macrophages, which are innate immune cells involved in both inflammation and tissue repair processes. *In vivo*, these cells exhibit a spectrum of phenotypes, ranging from pro-inflammatory (M1-like) to anti-inflammatory (M2-like), each having unique functional roles^41,42^. Recent work demonstrated that in bone marrow-derived macrophages (BMDM), lamins A/C are rapidly degraded upon pro-inflammatory stimulation, and lamin A/C-deficient macrophages exhibit increased expression of pro-inflammatory markers upon stimulation^30^. Nonetheless, the impact of pro-inflammatory polarization-induced lamin A/C degradation on nuclear mechanics in macrophages remains largely unexplored. Furthermore, previous studies were limited by the relatively short polarization durations of ≤6 hours and/or low-cell-number measurements (<10 cells) due to the use of low-throughput mechanical assays such as optical tweezers and acoustic force spectroscopy^43,44^. In contrast to M1-like macrophages, M2-like macrophages do not alter expression of lamins upon polarization^45,46^, leading us to focus our study on pro-inflammatory (M1-like) polarization and unpolarized macrophages.

To address this knowledge gap, we performed complementary physical measurements to determine the effect of pro-inflammatory stimulation on macrophage nuclear mechanics. We employed several advanced techniques targeting distinct mechanical regimes, including external force applications using micropipette aspiration and whole-cell migration through transwell assays, and intrinsic nuclear measurements using Brillouin microscopy, fluorescence lifetime imaging (FLIM), and chromatin tracking analysis to assess nuclear mechanics in macrophages. These approaches enabled a direct comparison of nuclear deformability, stiffness, and chromatin dynamics between unpolarized (M0) and pro-inflammatory (M1-like) macrophages in intact cells. Although pro-inflammatory macrophages substantially reduced the expression of nuclear lamins, we observed nuclear *stiffening* and *decreased* nuclear deformability compared to unpolarized macrophages. The pro-inflammatory macrophages were further characterized by a decrease in nuclear volume, redistribution of specific heterochromatin marks, and reduced intranuclear chromatin movement. These findings suggest that in primary macrophages, which have low lamin A/C levels compared to other cell types, lamins are not the primary determinant of nuclear mechanics, but that instead polarization-induced chromatin compaction drives nuclear stiffening and resistance to large nuclear deformations.

## Results

### Pro-inflammatory macrophages have decreased Lamin levels and altered nuclear morphology

Primary macrophages provide an excellent model for studying nuclear mechanics because they tend to more accurately represent physiological conditions of monocyte-derived macrophages *in vivo* compared to macrophage cell lines^47^. To generate primary macrophages, we isolated and froze bone marrow progenitor cells from the femurs and tibias of C57BL/6 mice. Prior to each experiment, we thawed the progenitor cells and treated the cells for 6 days with macrophage colony-stimulating factor (M-CSF) to induce macrophage differentiation (Supplementary Figure 1A). After 6 days of differentiation, macrophages exhibited around ∼96% cell viability (Supplementary Figure 1D, E) and significantly higher levels of F4/80 (Supplementary Figure 1F, G), a cell surface glycoprotein present on mature macrophages^48,49^, compared to an isotype control. Additionally, macrophages exhibited significantly higher levels of CD11b (Supplementary Figure 1H, I), a marker used to distinguish myeloid cells^50^. These observations indicate successful and robust differentiation of a highly pure macrophage population similar to previously established protocols^51^. We then polarized macrophages towards an M1-like, or pro-inflammatory, phenotype using stimulation with lipopolysaccharide (LPS) and interferon-gamma (IFNγ) (Fig. 1A), a well-established protocol to induce pro-inflammatory gene expression and cytokine secretion while mimicking the *in vivo* acute inflammatory response^41,42^. Henceforth, for simplicity, we refer to these cells as “M1” macrophages. As controls, we maintained a subset of macrophages from the original population that did not receive LPS or IFNγ and that represent a non-polarized phenotype, referred to as “M0” in this study (Fig. 1A).

**Figure 1.**
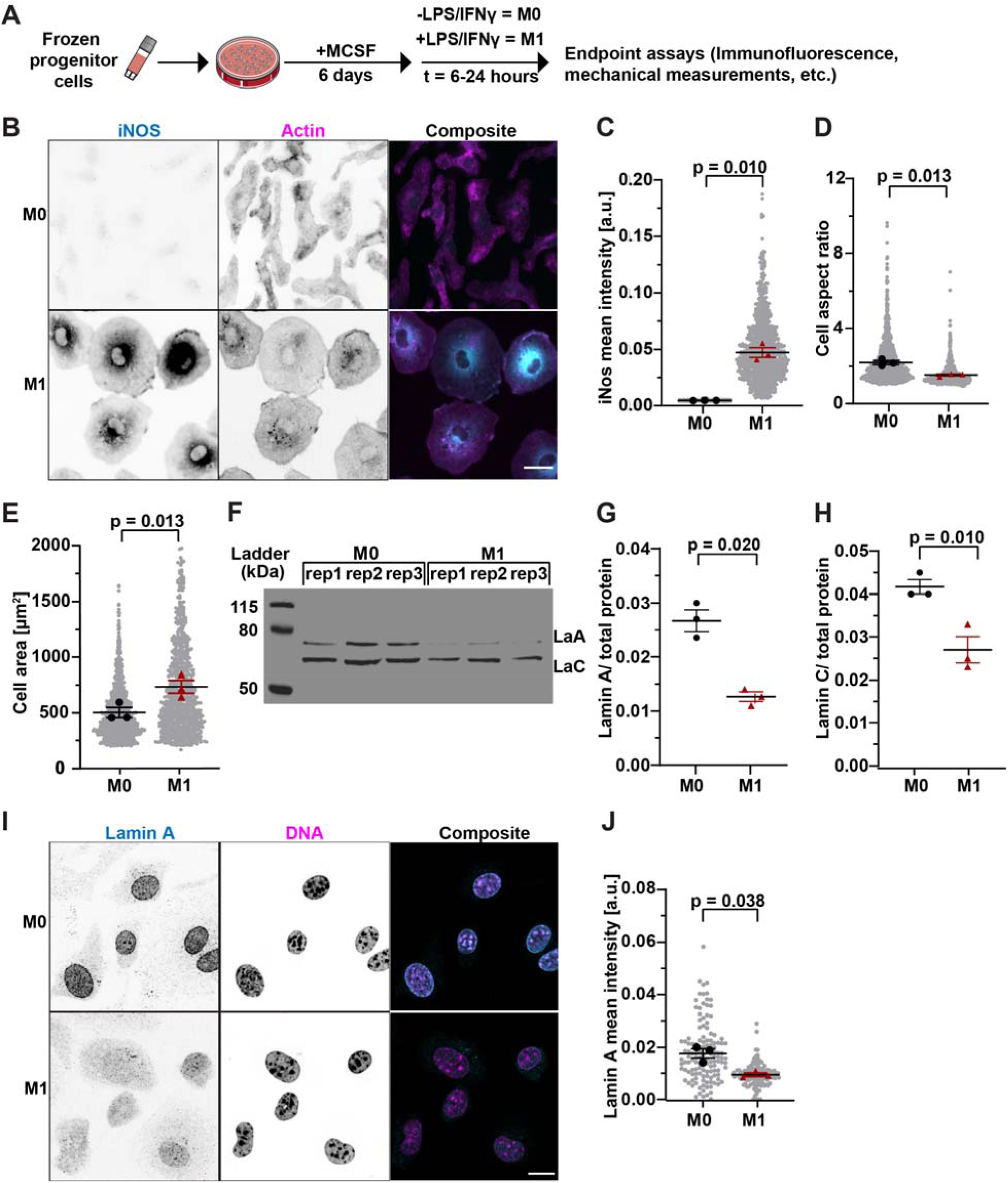
Primary bone-marrow derived macrophages have reduced lamin A/C levels upon pro-inflammatory stimulation. (**A**) Overview of macrophage differentiation and polarization timeline. (**B**) Representative images of iNOS expression in M0 or M1 macrophages polarized with LPS (100 ng/mL) and IFNγ (20 ng/mL). Scale bar = 20 µm. (**C**) Quantification of mean iNOS immunofluorescence intensity in M0 and M1 macrophages. (**D**) Quantification of the aspect ratios of M0 and M1 macrophages following polarization. The aspect ratios were calculated as major axis length/minor axis length. (**E**) Quantification of cell area of M0 and M1 macrophages. (C-E) *n* = 846-1195 cells per condition. (**F**) Representative immunoblot for lamins A/C of M0 and M1 macrophages. The blot used lysates from three independent experiments. (**G**) Corresponding quantification of lamin A levels in the immunoblot shown in panel F, normalized to total protein loading (Supplementary Figure 3) (**H**) Corresponding quantification of lamin C expression, based on immunoblot in F, normalized to total protein loading. (**I**) Representative maximum intensity projection images of cells immunofluorescently labeled for lamin A. Scale bar = 10 µm (**J**) Quantification of immunofluorescence images of cells stained for lamin A. *n* = 117-127 cells per condition. For all quantification involving individual cell measurements (C,D,E,K) data is shown as a line at the mean ± SEM, with replicate averages plotted in large black circles (M0) or red triangles (M1), and individual measurements shown as small grey circles. For immunoblot quantification (G, H) data is shown as a line at the mean ± SEM, with the replicate means plotted. Statistical analyses are based on paired *t*-tests using replicate means. Panel A icons provided by the Bioicons community from Servier (https://smart.servier.com/) is licensed under CC-BY 3.0.

Following treatment with pro-inflammatory stimuli, we immunofluorescently labeled macrophages with an antibody labeling inducible nitric oxide synthase (iNOS), a key marker of M1 macrophage polairzation^42^. M1 macrophages had significantly higher levels of iNOS expression compared to M0 macrophages (Fig. 1B, C). Consistent with previous reports of cell morphology changes in response to pro-inflammatory polarization^52,53^, M1 macrophages displayed altered cell morphology, including increased cell spreading as measured by a decrease in cell aspect ratio (Fig. 1D) and increased cell area (Fig. 1E) compared to unpolarized cells. These data confirm successful polarization of M1 macrophages by LPS and IFNγ stimulation.

To validate prior results^30^, we assessed levels of nuclear lamins following M1 polarization. As expected, M1 macrophages had significantly decreased levels of lamins A/C compared to M0 macrophages, based on both immunoblotting (Fig. 1F-H) and immunofluorescence labelling (Fig. 1I, J). In contrast, although lamin B1 levels showed a slight decrease, this change was not statistically significant (Supplementary Figure 2A, B, E, F). We also assessed the levels of additional nuclear envelope proteins, including lamin B receptor (LBR), as LBR levels correlate with nuclear envelope wrinkling and lobulation in neutrophils^54^ and LBR tethers heterochromatin to the periphery of the nucleus^29,55–57^, and emerin, which contributes to nuclear shape and mechanics^58,59^. LBR expression was slightly, but statistically significantly decreased following polarization (Supplementary Figure 2A, C), indicating that the nuclear envelope composition changes beyond lamins A/C. In contrast, emerin expression did not change upon M1 polarization (Supplementary Figure 2G, H).

To directly compare the levels of lamin A/C expression in the bone marrow-derived macrophages to those in other cell types, we assessed lamin A/C expression in M0 macrophages, 4T1 mouse mammary carcinoma cells, which have relatively soft nuclei and low lamin A/C levels compared to other breast epithelial cells^60^, and mouse embryonic fibroblasts (MEFs), which have relatively high lamin A/C levels and that have been extensively characterized for their nuclear mechanics^31,32,61^. Lamin A levels were significantly lower in M0 macrophages compared to both 4T1 cells and MEFs (Supplementary Figure 2J, K). In contrast, lamin C levels in M0 macrophages were slightly lower than in MEFs and significantly lower than in 4T1 cells (Supplementary Figure 2J, L).

Since changes to the nuclear lamina can directly alter nuclear volume and shape^37^, we assessed if M1 macrophages had altered nuclear morphology compared to M0 controls. Using high-resolution 3D confocal imaging, we observed a significant decrease in nuclear volume, area, height, and circularity relative to unpolarized M0 macrophages (Fig. 2A-E). We also quantified the proportion of cells with nuclear invaginations, tunnels, and circumferential wrinkling (Fig. 2F-H). Here, nuclear invaginations were defined as indentations in the nuclear envelope, tunnels were identified as nuclear envelope features that transverse the entire nucleus, and circumferential wrinkling was identified as wrinkling around the periphery of the nucleus. While we did not observe any statistically significant differences in the proportion of nuclei that contained nuclear invaginations (Fig. 2I) or tunnels (Fig. 2J) between M0 and M1 macrophages, M1 macrophages had a significant increase in the proportion of nuclei with circumferential nuclear envelope wrinkling compared to M0 controls (Fig. 2K). Furthermore, M1 macrophages had a significantly higher percentage of excess nuclear perimeter (Fig. 2L), defined as the difference between the nuclear perimeter and the perimeter of a circle with the equivalent area. These findings show that following polarization, M1 nuclei decrease in volume and exhibit increased excess nuclear membrane surface, as measured by nuclear perimeter, thus leading to nuclear envelope wrinkling.

**Figure 2.**
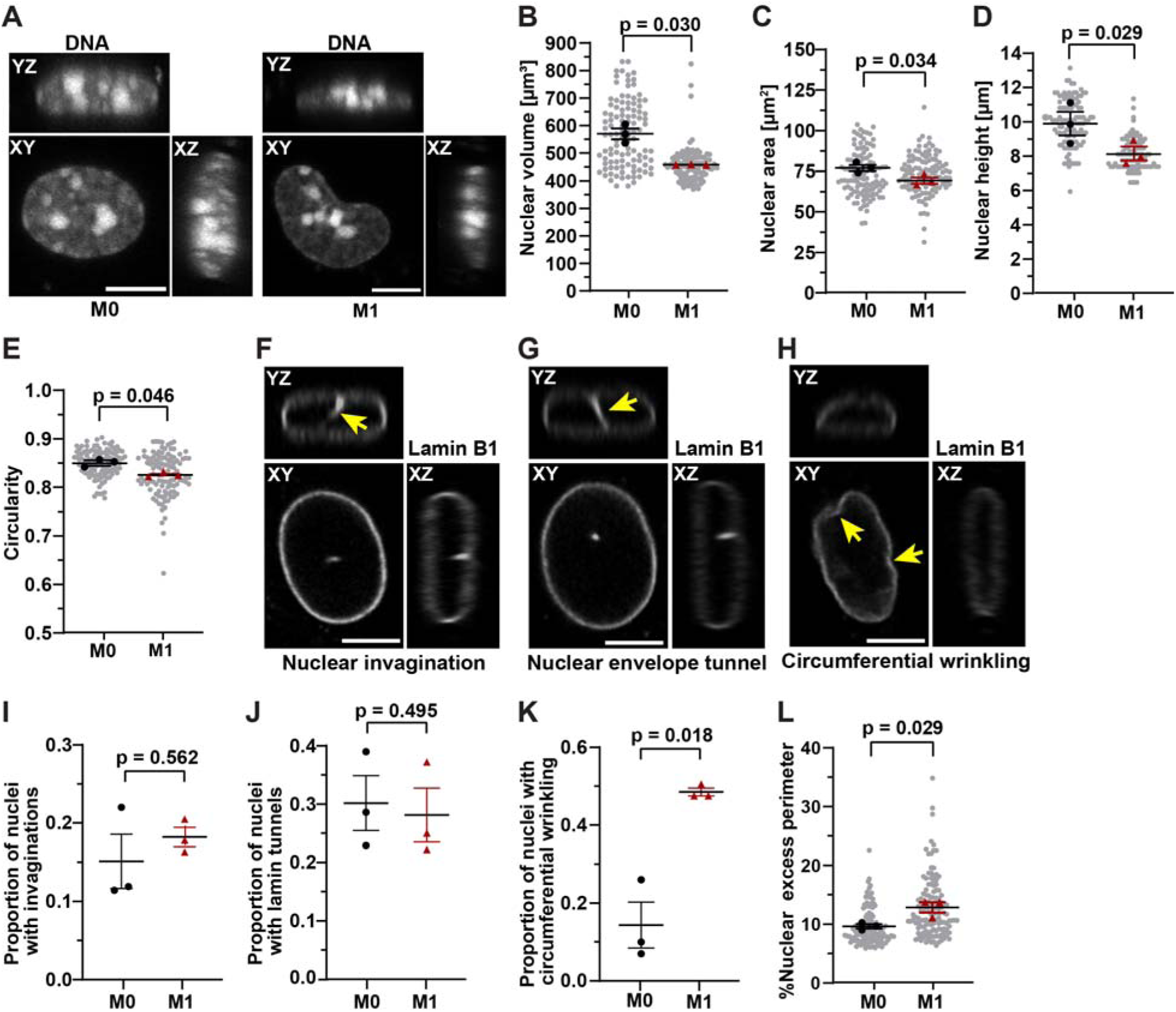
Pro-inflammatory macrophages have decreased nuclear volume and altered nuclear morphology. (**A**) Representative orthogonal view projections of 3D image z-stacks of M0 and M1 macrophages stained with DAPI to reveal nuclear morphology. The images show XY projection and XZ, YZ orthogonal views. Scale bar = 5 µm. (**B-E**) Quantification of nuclear volume (**B**), nuclear area (**C**), nuclear height (**D**), and circularity (**E**) of M0 and M1 macrophages. *n* = 107-109 cells per condition from three independent experiments. (**F-H**) Representative orthogonal view projections of 3D image z-stacks of a nucleus with nuclear envelope invaginations (**F**), nuclear envelope tunnels (**G**), and nuclear circumferential wrinkling (**H**), marked by yellow arrow. The nuclear envelope was visualized based on immunofluorescence labeling for lamin B1. Scale bar = 5 µm. (**I-K**) Quantification of nuclear features including proportion of nuclei with nuclear envelope invaginations (**I**) lamin tunnels (**J**) and proportion of nuclei with only circumferential wrinkling (**K**), based on lamin B1 immunofluorescence analysis. (**L**) Quantification of the excess perimeter of the nuclear envelope (expressed as percentage) of M0 and M1 macrophages based on lamin B1 staining. For all quantifications involving individual cell measurements (B-E, L) data are shown as a horizontal line at the mean ± SEM, with averages from individual experimental replicates plotted in large black circles (M0) or red triangles (M1), and individual nuclear measurements shown as small grey circles. For nuclear wrinkling quantification (I, J) data is shown as a horizontal line at the mean ± SEM, with the replicate means plotted. *n* = 83-135 cells per condition. Statistical analyses are based on paired *t*-tests using replicate means.

Because we observed a loss of nuclear volume in M1 macrophages, we assessed the DNA content and cell-cycle distribution of M0 and M1 macrophages, given that differences in macrophage cell cycle states are known to arise from pro-inflammatory stimulation^62^, and cell cycle progression influences nuclear DNA content. We performed cell cycle analysis using propidium iodide staining to quantify DNA content and identify cell cycle phase in M0 and M1 macrophages (Supplementary Figure 4A, B). M1 macrophages had a significant higher percentage of cells in G0/G1 phase, and also a slight increase of cells in G2/M phases, whereas the fraction of M1 macrophages in S phase was significantly reduced compared to M0 controls (Supplementary Figure 4C), consistent with previous studies^63^. Total DNA content was not statistically significant between M0 and M1 macrophages, despite the differences in nuclear volume, suggesting that M1 cells have a higher DNA density, indicating chromatin compaction. Additionally, the increased fraction of M1 macrophages in G0/G1 and the lower fraction of these cells in S phase suggest that M1 cells have less replicating DNA, which might allow for more chromatin compaction.

In addition, we assessed if alterations in nuclear volume affect large scale chromatin organization, such as the size and distribution of heterochromatic chromocenters^64,65^. To answer this, we performed image analysis on the M0 and M1 macrophage chromocenters, identified based on punctae with high DAPI intensity. We did not observe any differences between the number, area, or distribution of chromocenters between M0 and M1 macrophages (Supplementary Figure 4 D-I). Since the ∼20% loss in nuclear volume we observed in M1 macrophages corresponds to only a ∼7% reduction in nuclear radius, it is possible that subtle changes in large-scale chromatin distribution occur during polarization but are too small to detect in our assays.

### Pro-inflammatory polarization leads to decreased nuclear deformability in macrophages

These differences in nuclear morphology between M0 and M1 macrophages, in particular the loss of nuclear volume and nuclear circumferential wrinkling, along with the decrease in lamin A/C levels prompted us to further elucidate differences in nuclear mechanics between polarized and unpolarized macrophages. Because M1 macrophages had decreased levels of Lamin A/C, we hypothesized that these cells would have more deformable nuclei compared to M0 macrophages. To test this hypothesis, we applied a recently developed microfluidic micropipette aspiration assay^66^ to impose large nuclear deformations to M1 and M0 macrophages (Fig. 3A). The micropipette aspiration assay allows for high-throughput single-cell quantification of nuclear deformations using time lapse imaging of nuclei stained with a live-cell DNA dye and an analysis pipeline to quantify protrusion length of individual nuclei into the micropipette channels^66^. Because a previous study found that *Lmna* mRNA levels start to decline within 3 hours of pro-inflammatory treatment^30^, and protein levels are expected to lag changes in mRNA levels by a few hours, we performed micropipette aspiration on M1 macrophages at 6 and 24 hours of polarization and compared the nuclear deformability to that of M0 control macrophages collected at the same time points (Fig. 3B). Counter to our hypothesis, M1 macrophages had a trend (*p* = 0.101) towards a *decrease* in nuclear deformability at 6 hours of polarization compared to the M0 controls, and nuclear deformability decreased significantly after 24 hours of polarization (Fig. 3C-E). We performed statistical analysis comparing the nuclear protrusion lengths at multiple time points, which revealed a significant decrease in nuclear protrusion length of M1 macrophages compared to M0 macrophages at 60 and 190 seconds of aspiration, and a slight but not quite statistically significant difference (p = 0.076) at 140 seconds (Supplemental Fig. 5 A-C). These results reveal an unexpected finding, in which M1 macrophages have *less* deformable nuclei compared to M0 macrophages, despite their decreased lamin A/C levels, indicating that other factors must contribute to the nuclear stiffening in the M1 cells.

**Figure 3.**
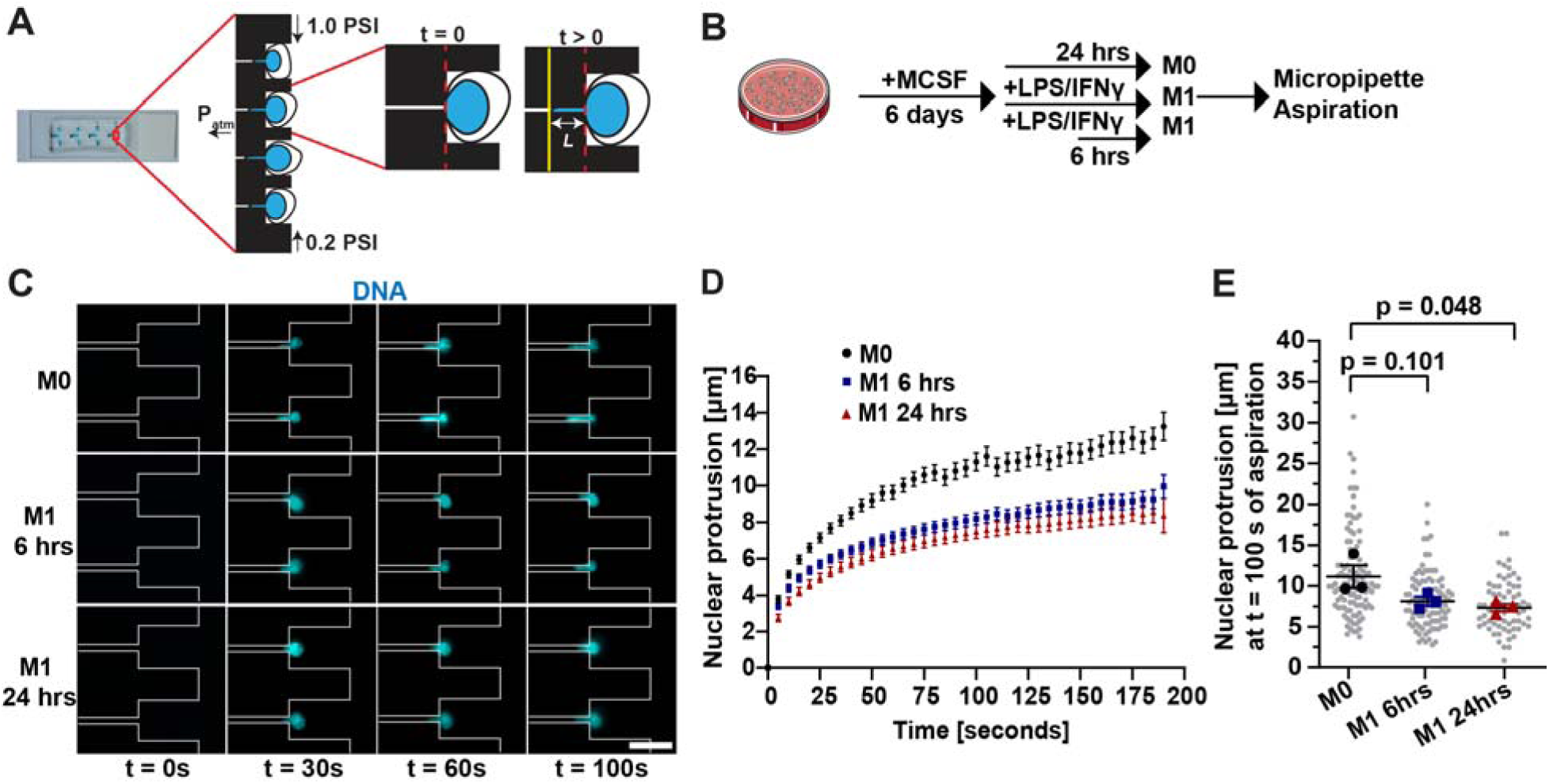
Pro-inflammatory polarization leads to decreased nuclear deformability in macrophages. (**A**) Schematic of microfluidic micropipette aspiration device modified, adapted from ^32^. (**B**) Overview of macrophage differentiation and polarization timeline for micropipette aspiration experiments. (**C**) Representative images of nuclear protrusions of M0 and M1 macrophages during micropipette aspiration. Boundaries of the micropipette device channels are shown in grey. Scale bar =20 µm. (**D**) Quantification of nuclear protrusion length over time. Plots show mean nuclear protrusion length ± SEM at each timepoint from three independent replicates. (**E**) Nuclear protrusion length measured at 100 seconds after the start of nuclear aspiration. *n* = 68-109 cells per condition. Data shown as mean ± SEM, with replicate averages plotted in large black circles (M0), blue squared (M1 6 hrs), or red triangles (M1 24 hrs), and individual measurements shown as small grey circles. Statistical analysis is based on a one-way ANOVA with Dunnett’s multiple comparison test using replicate means. Panel B icon provided by the Bioicons community from Servier (https://smart.servier.com/) is licensed under CC-BY 3.0.

Since pro-inflammatory signaling resulted in changes in the aspect ratio and spreading of M1 macrophages (Fig. 1D, E), and prior studies found that polarization leads to changes in the actin cytoskeleton^67^, we considered whether the observed difference in nuclear stiffness may be caused by altered actin organization between M1 and M0 macrophages. To address this question, we performed microfluidic micropipette aspiration on M0 and M1 macrophages treated with cytochalasin D to inhibit actin polymerization (Fig. 4A, B), thus allowing us to distinguish between nucleus-intrinsic stiffness versus apparent stiffening effects mediated through the actin cytoskeleton. These experiments were conducted after 24 hours of polarization (Fig. 4A), when M1 macrophages show a significant reduction in nuclear deformability (Fig. 3D, E; Fig. 4C). Treatment with Cytochalasin D resulted in a general increase in nuclear deformability in both M0 and M1 macrophages (Fig. 4C, D). However, even with cytochalasin D treatment, M1 macrophages had significantly less deformable nuclei compared to M0 macrophages (Fig. 4 B, E). In addition, whereas many cytochalasin D treated M0 cells were able to completely pass through the aspiration channel during the acquisition period, indicating their high deformability, significantly fewer cytochalasin D treated M1 macrophages transited the microchannels (Fig. 4F), further validating that M1 macrophages have stiffer nuclei with reduced deformability, and that the increased stiffness of M1 macrophages observed in our assays is independent of differences in actin organization.

**Figure 4.**
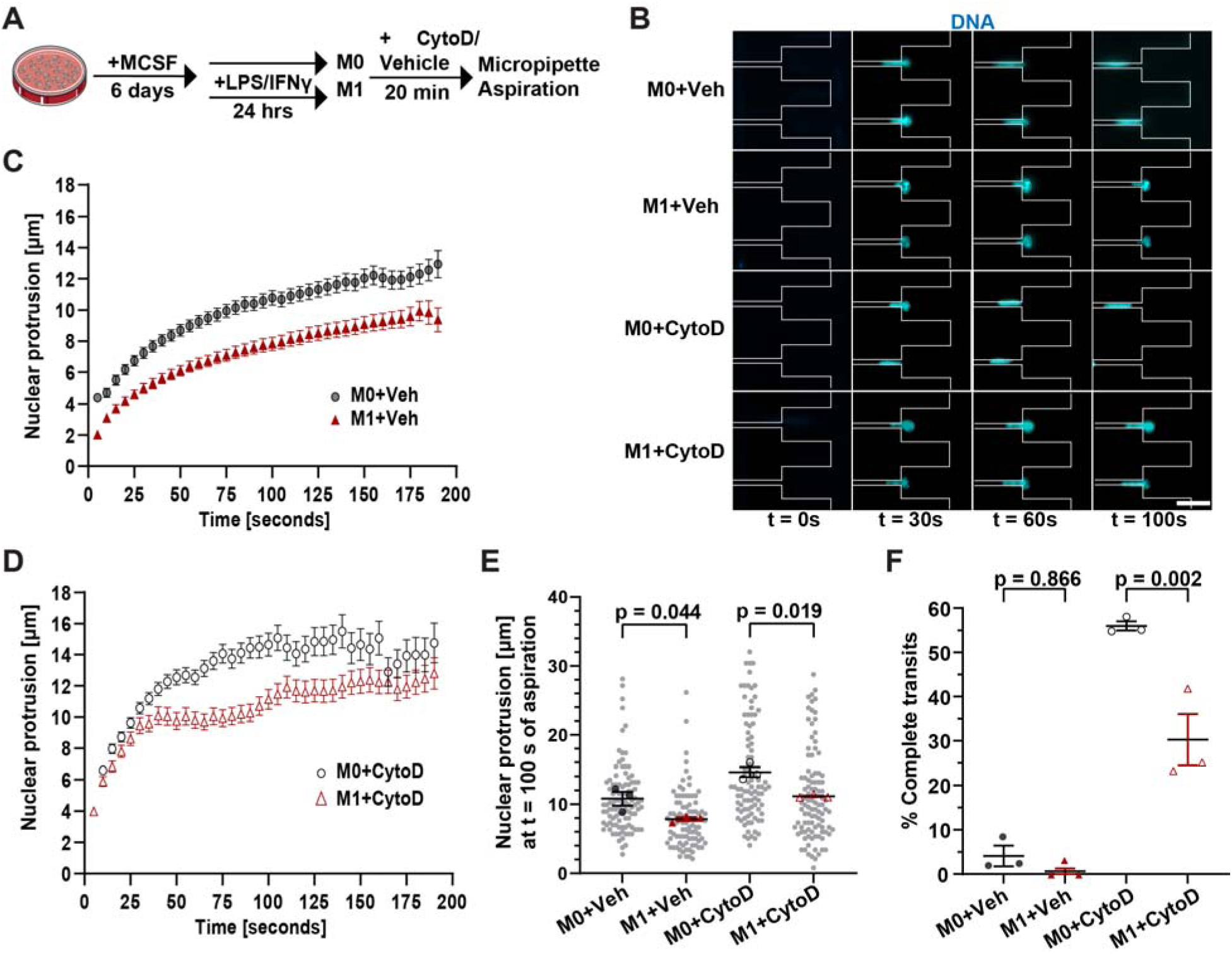
Increased nuclear stiffness in M1 macrophages is not dependent on the actin cytoskeleton. (**A**) Overview of macrophage differentiation and polarization timeline for micropipette aspiration experiments including cytochalasin D treatment. (**B**) Representative images of nuclear protrusions of M0 and M1 macrophages with and without cytochalasin D treatment during micropipette aspiration. Boundaries of the micropipette device channels are shown in grey. Scale bar =20 µm. (**C**) Quantification of nuclear protrusion length over time for M0 and M1 macrophages + vehicle control. (**D**) Quantification of nuclear protrusion length over time for M0 and M1 macrophages treated with Cytochalasin D (CytoD). (**E**) Nuclear protrusion length at 100 seconds after the start of nuclear aspiration. Plots (D, E) show mean nuclear protrusion length ± SEM at each time point from three independent replicates. *n* = 92-101 cells per condition from three independent replicates. (**F**) Quantification of the proportion of nuclei that completely transit through the micropipette channel for each condition. For quantification involving individual cell measurements (E) data is shown as a line at the mean ± SEM, with replicate averages plotted in large black circles (M0) or red triangles (M1), and individual measurements shown as small grey circles. For complete transit data (F) data is shown as a line at the mean ± SEM, with the replicate means plotted. Statistical analysis is based on one-way ANOVA with Tukey’s multiple comparison test using replicate means. Panel A icon provided by the Bioicons community from Servier (https://smart.servier.com/) is licensed under CC-BY 3.0.

As an orthogonal measure of nuclear stiffness, we performed Brillouin microscopy, a noninvasive label-free technique that uses light scattering to probe the mechanical properties of materials by measuring the frequency shift caused by the interaction of light with acoustic waves in the sample, known as the Brillouin frequency shift (BFS)^68,69^. This frequency shift can be quantified and provides a characterization of the mechanical properties of subcellular structures in living cells^61,70^. Segmenting the cell nucleus, based on a cell permeable fluorescent DNA stain (Supplementary Figure 5D), enabled us to assess nuclear stiffness in live M0 and M1 macrophages. M1 macrophages had an increased Brillouin frequency shift compared to M0 macrophages (Supplementary Figure 5D, E), indicating that M1 macrophages have stiffer nuclei than M0 macrophages, consistent with our micropipette aspiration assays. Collectively, our micropipette and Brillouin data suggest that pro-inflammatory macrophage polarization results in stiffer and less deformable nuclei, and that this change in nuclear mechanics cannot be explained by changes in lamin levels or actin organization associated with polarization.

### Pro-inflammatory macrophages have altered H3K9me3 localization, increased chromatin compaction, and altered chromatin dynamics

In addition to lamins, chromatin organization can also modulate nuclear deformability^15,22,37^, and cell differentiation and fate decisions are often associated with changes in chromatin organization^71–73^. Furthermore, lamins and LBR contribute to chromatin localization by tethering heterochromatin to the periphery of the nucleus via lamina associated domains (LADs)^29,55–57^. Thus, we considered whether M1 macrophage polarization results in changes to chromatin compaction and mobility. We immunofluorescently labeled M0 and M1 macrophages with antibodies recognizing trimethylated histone H3K9 (H3K9me3) and H3K27 (H3K27me3), markers of constitutive and facultative heterochromatin, respectively. We also labeled cells using an antibody recognizing Histone-3 (H3), as the ratio of heterochromatin-specific stains to total H3 allows for cell-by-cell normalization to account for differences in antibody penetrance or sample permeabilization. The ratio of H3K9me3 to H3 nuclear intensity was unchanged following polarization (Supplementary Figure 6A-D) but M1 macrophages exhibited a distinct change in subnuclear distribution of the H3K9me3 signal. Intensity profiles measuring H3K9me3 signal across a line drawn through the midplane of the nucleus revealed that M1 macrophages had significantly less H3K9me3 at the nuclear periphery compared to M0 macrophages (Fig. 5A, B) and a decreased ratio of peripheral to nucleoplasmic H3K9me3 intensity (Fig. 5C). In contrast to the constitutive heterochromatin mark H3K9me3, the levels and subnuclear distribution of the facultative heterochromatin mark H3k27me3 remained unchanged following M1 polarization (Supplementary Figure 6E-H). These results suggest that M1 polarization induces a change in specific chromatin organization, with constitutive heterochromatin marks lost from the nuclear periphery, perhaps as a consequence of reduced lamin A/C levels or to facilitate the expression of pro-inflammatory genes. Indeed, previous studies have shown H3k9me3 present at inflammatory gene loci decrease after LPS stimulation to allow for transcription of pro-inflammatory genes^74–76^; however. genome wide changes in H3K9me3 or altered localization of H3k9me3 in pro-inflammatory macrophages have not been reported previously.

**Figure 5.**
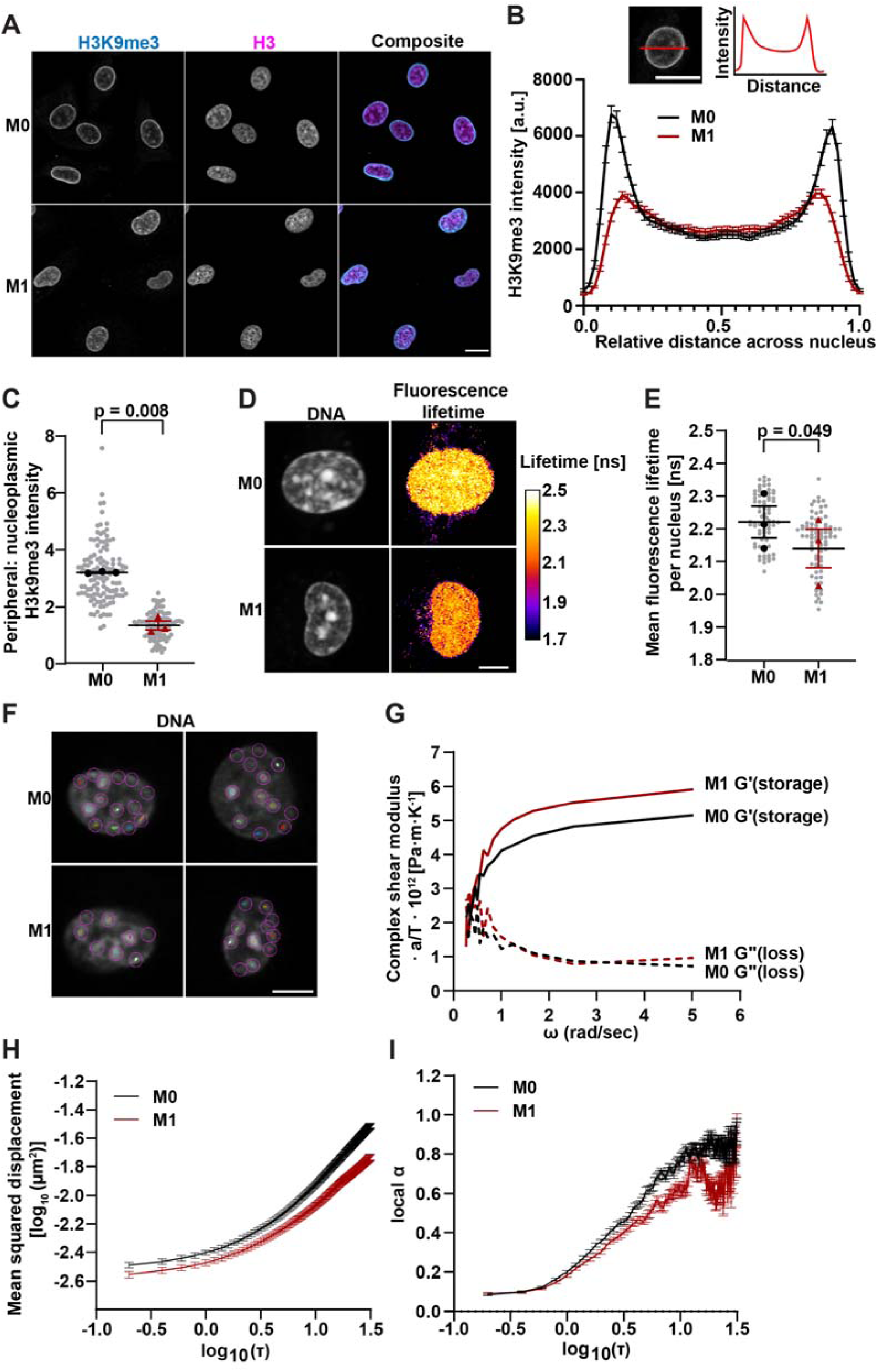
Pro-inflammatory macrophages have altered H3K9me3 localization, chromatin compaction and chromatin dynamics. (**A**) Representative single confocal image slices of cells immunofluorescently labeled for H3K9m3 and Histone-3 (H3). Scale bar = 10 µm. (**B**) Quantification of H3K9me3 intensity profiles along a line through the midplane of the nucleus of M0 (black line) and M1 (red line) macrophages. (**C**) Quantification of the ratio of H3k9me3 at the nuclear periphery to H3k9me3 in the nucleoplasm as shown in (B). *n* = 94-117 cells per condition. (**D**) Representative images of FLIM data showing Hoechst staining and weighted average fluorescence lifetime intensity [ns] per pixel. Scale bar = 10 µm. (**E**) Quantification of mean fluorescence lifetime per nucleus of M0 and M1 macrophages. *n* = 73-82 cells per condition. (**F**) Representative images of M0 and M1 nuclei stained with Hoechst 33342 with chromatin puncta tracks identified. Scale bar = 5 µm. (**G**) Quantification of the complex shear modulus, represented by the storage modulus (G’) and loss modulus (G’’) for M0 and M1 macrophage nuclei, shown as G’a/T and G’’a/T, where *a* is the chromatin foci size and *T* is the effective temperature. Because both *a* and *T* are unknown, G’a/T and G’’a/T were reported rather than absolute moduli *G’* and *G’*’. (H) Log-log plot of ensemble-time-averaged MSD vs. time lag (τ) of M0 and M1 nuclei. (**I**) Log-log plot of local α values vs. time lag (τ) for M0 and M1 nuclei. For all quantification involving individual cell measurements (C,E) data is shown as a line at the mean ± SEM, with replicate averages plotted in large black circles (M0) or red triangles (M1), and individual measurements shown as small grey circles. For line plots (B,H,I) data is plotted as mean ± SEM. Statistical analyses are based on paired *t*-tests using replicate means.

Since M1 macrophages had decreased nuclear volume (Fig. 2B) and altered subnuclear heterochromatin distribution (Fig. 4A-C), we hypothesized that M1 polarization leads to chromatin condensation that could explain the increased stiffness observed in the micropipette aspiration and Brillouin microscopy assays. To quantify chromatin compaction, we performed fluorescence lifetime imaging microscopy (FLIM) on macrophages stained with Hoechst, as the fluorescence lifetime of DNA-bound Hoechst is inversely related to the local compactness of chromatin^77^. M1 macrophages had a significantly lower fluorescence lifetime of Hoechst than M0 cells (Fig. 5D, E), indicating that pro-inflammatory polarization promotes chromatin condensation in M1 macrophages.

To validate our findings of increased chromatin condensation in M1 macrophages, we assessed chromatin dynamics in M0 and M1 macrophages by performing live cell imaging of macrophage nuclei followed by single particle tracking methods to quantify chromatin motion over short time scales (Fig. 5F). Using a custom MATLAB script implementing the generalized Stokes-Einstein equation, we observed a higher elastic storage modulus (*G*’) of M1 macrophage nuclei compared to M0, indicating that M1 nuclei are stiffer compared to M0 (Fig. 5G). In contrast, the loss modulus (*G*”) did not differ significantly between M1 and M0 macrophage nuclei (Fig. 5G), suggesting similar viscous properties. In addition, we calculated and plotted the ensemble-time-averaged mean squared displacement (MSD) as a function of time lag (τ) on a log-log scale (Fig. 5H) for chromatin tracks, where we observed a decreased chromatin motion in M1 nuclei relative to M0. To further examine the viscoelastic characteristics of chromatin at different time scales, we applied a sliding window linear model fit to determine the local anomalous exponent α, derived from the slope of the MSD-r relationship on the log-log plot. Higher α values indicate a less stiff environment due to less molecular crowding, whereas lower α values indicate a stiffer environment due to increased molecular crowding^38,78,79^. Plotting local α as a function of log(τ) revealed consistently lower α values in M1 nuclei compared to M0 nuclei (Fig. 5I). Taken together, these results indicate that M1 macrophage nuclei exhibit a stiffer elastic modulus compared to unpolarized M0 macrophages, driven by altered chromatin organization and dynamics in the nuclear interior.

Nuclear mechanics play a key role in confined migration, as cells must generate intracellular forces to extensively deform their nuclei to pass through small physical barriers^6,7^. Previous studies have shown that cells with less deformable or less stiff nuclei as a result of decreased lamin A/C levels migrate more efficiently through confined spaces^8,60^. Because M1 macrophages had stiffer nuclei than M0 controls in our experiments, we predicted that M1 macrophages are less efficient at migrating through confined spaces. To test this prediction, we compared the ability of M0 and M1 macrophages to migrate through transwell membranes with different pore sizes, ranging from 3 µm in diameter to 8 µm in diameter, representing a spectrum of local confinement. We assessed macrophage migratory ability by quantifying the number of cells that completed migration through transwell membranes after 6 and 24 hours (Fig. 6A). Although we did not observe any statistically significant difference between M0 and M1 migration through 3 µm pores after 6 hours, M1 macrophages were significantly less migratory at 24 hours, (Fig. 6B-E). Notably though, M1 macrophages also exhibited reduced migration through the larger 5 µm and 8 µm pores (Fig. 6B-E), which do not require extensive nuclear deformation, indicating that M1 macrophages are overall less efficient at migrating, regardless of the pore size. Because *in vivo* macrophages often migrate in response to a chemokine stimulus^80^, we additionally performed migration studies with the addition of fMLP (*N*-Formylmethionyl-leucyl-phenylalanine), a chemoattractant that promotes macrophage migration^81^, and observed similar results in which M1 macrophages were overall less migratory than M0 macrophages through all pore sizes, irrespective of chemokine stimulation (Supplementary Figure 7A, B). These data confirm that although M1 macrophages with stiffer nuclei exhibit decreased migratory ability compared to M0 macrophages—both in the absence and presence of chemokine stimulation—other factors beyond nuclear deformability contribute and likely dominate the migratory ability of these cells. Previous studies have identified that M1 macrophages exhibit less random migration on 2D substrates and less infiltration into 3D matrices compared to other macrophage phenotypes^43,82–84^. Additionally, pro-inflammatory polarization promotes increased cell adhesion^82,85,86^ and increased cell-cell clustering^45^, both of which could be the cause of decreased migratory ability in M1 macrophages. Other factors such as cytoskeletal dynamics and cell-substrate interactions likely contribute further to the reduced migratory potential of these cells^87,88^. Our work identifies increased nuclear stiffness as another possible contributor to the decreased migratory ability of M1 macrophages in confined environments, which might be partially offset by the reduced nuclear size of these cells. Future work is needed to evaluate the relative contributions of these factors in specific migration contexts.

**Figure 6.**
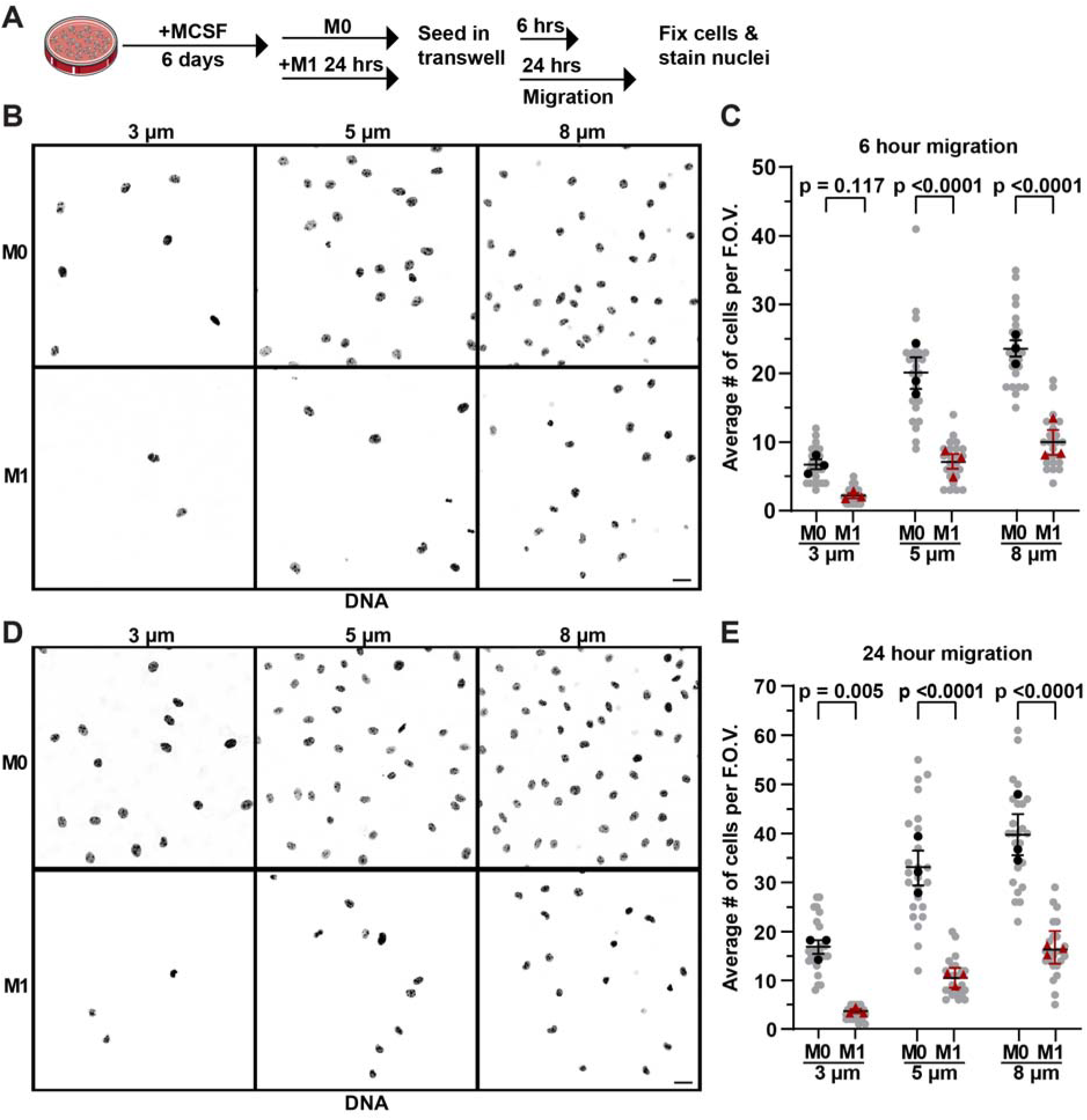
M1 macrophages exhibit reduced transwell migration in both confined and un-confined environments. (**A**) Overview of macrophage differentiation and polarization timeline for transwell migration experiments. (**B**) Representative images of macrophage nuclei stained with DAPI after 6 hours of migration through transwell membranes with either 3 µm, 5 µm, or 8 µm diameter pores. Scale bar = 20 µm. (**C**) Quantification of the number of cells that completed migration through transwell membranes by 6 hours. (**D**) Representative images of macrophage nuclei stained with DAPI after 24 hours of migration through transwell membranes with 3 µm, 5 µm, or 8 µm diameter pores. Scale bar = 20 µm. (**E**) Quantification of the number of cells that completed migration through transwell membranes by 24 hours. For each replicate, 4 fields of view were taken from 2 transwell devices, totaling 8 fields of view per condition per replicate. Data is plotted as a line at the mean ± SEM, with replicate averages plotted in large black circles (M0) or red triangles (M1), and individual measurements shown as small grey circles. Statistical analysis are based on two-way ANOVA with Sidak’s multiple comparison test using replicate means. Panel A icon provided by the Bioicons community from Servier (https://smart.servier.com/) is licensed under CC-BY 3.0.

## Discussion

In this study, we provide a comprehensive characterization of nuclear mechanics in pro-inflammatory murine macrophages. Surprisingly, while pro-inflammatory macrophages have reduced protein levels of lamins A/C, they exhibit less deformable, i.e., stiffer nuclei. This contrasts with reports from other cell types, where reduced lamin A/C levels typically correspond to increased nuclear deformability and decreased nuclear stiffness. Notably, the increased nuclear stiffness of pro-inflammatory macrophages was not due to differences in their actin cytoskeletons, as nuclei of M1 cells remained significantly stiffer than M0 cells following disruption of actin networks with cytochalasin D. Our findings suggest that in primary, pro-inflammatory macrophages, factors other than lamins or actin primarily determine nuclear mechanics.

Pro-inflammatory M1 macrophages had decreased nuclear volume and increased nuclear envelope wrinkling (Fig. 2B, K), while maintaining similar total amounts of DNA per nucleus (Supplementary Figure 4D), suggesting that inflammatory signals drive nuclear shrinkage and chromatin compaction. Since the proportion of nuclei with invaginations or tunnels did not correlate with the increased nuclear stiffness in M1 macrophages, we infer that nuclear invaginations/tunnels are not the major drivers for the difference in nuclear stiffness between M0 and M1 macrophages. Moreover, FLIM measurements showed that M1 macrophage chromatin exhibited shorter lifetimes (Fig. 5D, E), consistent with increased chromatin condensation compared to M0 macrophages, which could explain the increase in nuclear stiffness. Brillouin microscopy confirmed that the increased nuclear stiffness of M1 macrophages was associated with a stiffer nuclear interior (Suppl. Fig. 4D, E), as Brillouin measures the stiffness of chromatin through the Brillouin frequency shift of scattered light^89–91^. Tracking chromatin dynamics in live cells similarly indicated increased nuclear stiffness (Fig. 5F-I), and pro-inflammatory macrophages had redistribution of the heterochromatin mark H3K9me3 from the nuclear periphery to the nuclear interior (Fig. 5B, C). We envision that the combination of these effects, including nuclear volume loss, chromatin condensation, and heterochromatin reorganization, are sufficient to stiffen the nuclei of pro-inflammatory macrophages despite the concomitant reduction in lamins A/C expression. This interpretation is also supported by the fact that macrophages exhibit much lower levels of lamin A/C than other cell types, including some with already very deformable nuclei, such as 4T1 cells (Supplementary Figure 2J-L). Therefore, further reducing lamin A/C levels upon pro-inflammatory stimulation in macrophages is unlikely to further soften the nucleus, whereas increasing chromatin compaction and crowding by reducing nuclear volume during polarization is likely driving the increased nuclear stiffness in M1-like macrophages.

Macrophages are key players in many disease pathologies including cancers, chronic inflammatory and autoimmune diseases, cardiovascular disease, and others^92–94^. In both innate immune responses and disease pathologies, macrophages are subject to changing physical environments with altered local forces acting on the cells, but it remains to be determined whether increased nuclear stiffness confers functional consequences to pro-inflammatory macrophages. Nuclear stiffening could alter mechanosensitive signaling pathways and enhance the pro-inflammatory response, or nuclear stiffening could be a passive side effect of chromatin compaction and changes to gene expression as cells activate pro-inflammatory pathways. Future work is needed to determine whether the increased nuclear stiffness of M1 macrophages contributes to their ability to release pro-inflammatory cytokines, generate reactive oxygen species, or perform phagocytosis. Since phagocytosis is heavily actin dependent, a stiffer nucleus might act as a more stable anchor for the generation of intracellular cytoskeletal forces to allow macrophages to perform phagocytosis more efficiently.

Another important aspect of macrophage nuclear mechanobiology is the ability of these cells to migrate through tissue architecture as they are recruited to sites of infection^95,96^. In many solid tumor cancers, macrophages make up a large portion of the tumor by mass^97^, and cells within the tumor microenvironment experience compressive stresses^98^. This raises the possibility that increased nuclear stiffness following polarization might be a mechanism to protect the genome from damage in mechanically stressful environments. Additionally, it is possible that once a macrophage becomes polarized, it no longer requires a deformable nucleus, as the advantages of nuclear deformability for migrating through confined spaces are diminished as polarized macrophages lose migratory capacity.

Our study has some limitations. First, primary BMDMs are not conducive to *ex vivo* genetic manipulation without inducing polarization, cell death, or degradation of the insertion target^99,100^. Our experiments were therefore limited to live cell stains and immunofluorescence labeling in fixed samples. Although macrophage cell lines exist that are amenable to genetic modification^101,102^, the commonly used RAW 264.7 cell line does not degrade lamins A/C following inflammatory stimuli^30^, and previous work using peritoneal elicited macrophages did not observe decreased lamin A/C after treatment with LPS^44^, which contrasts with BMDMs^30^ and indicates cell-type specific differences that would confound these analyses. Additionally, although previous studies reported nuclear *softening* of isolated THP-1 macrophage nuclei after short-term (<6 h) LPS treatment, these studies also observed LPS-induced nuclear volume loss in THP-1 cells, human-derived macrophages, and peritoneal macrophages^44^. These results suggest that nuclear volume loss is a common response following inflammatory stimulation, but also highlight cell type specific differences on nuclear stiffness in response to polarization, at least at short time-scales. Finally, we chose to study nuclear mechanics in intact cells to avoid potential damage to the nuclear envelope during nuclear isolation and altering nuclear stiffness due to changes in multi-valent ion concentrations. Although this means that the cytoplasmic cytoskeleton may contribute to the mechanical measurements, and we thus report ‘effective’ nuclear stiffness, we believe that the importance of maintaining a physiological nuclear environment justifies this choice. Lastly, even though nuclear deformability differences persist when inhibiting actin polymerization, other cytoskeletal components may contribute to nuclear mechanics, including microtubules and cytoplasmic intermediate filaments.

The detailed molecular mechanism driving chromatin condensation and nuclear stiffening in the polarized macrophage remains to be fully determined, and it is currently unknown whether nuclear volume loss is the driver or result of chromatin condensation. Future studies should aim to elucidate the biophysical mechanisms and molecular players in nuclear stiffness of macrophages. Such studies, however, would likely require primary macrophages from genetically engineered mouse models or *ex vivo* genetic modifications, such as modulation of lamin A/C expression, or the introduction of fluorescent markers to study chromatin dynamics and compaction in more detail. Additionally, open questions exist as to whether the compaction of the nuclear volume alone is sufficient to explain nuclear stiffening in pro-inflammatory macrophages, or if other factors such as altered interactions between chromatin and the nuclear envelope further contribute to changes in nuclear mechanics of these cells. It is also unclear whether the change in nuclear stiffness during polarization is the consequence of altered chromatin organization required for altered gene expression associated with pro-inflammatory polarization. Additionally, future work is needed to investigate the plasticity of nuclear stiffness changes following M1 polarization. Careful consideration is needed to deduce whether molecular changes during pro-inflammatory stimulation directly alter nuclear mechanics or modulate other pathways capable of modulating nuclear stiffening during polarization, making it challenging to answer these mechanistic questions. For example, perturbing changes in chromatin organization associated with M1 polarization would interfere with polarization itself, precluding us from delineating between these scenarios.

Taken together, our findings have broad relevance for macrophage cell biology and mechanobiology, as macrophages are key players in many physiological processes and disease pathologies. The present study expands our understanding of macrophage nuclear mechanics by showing that in these cells, chromatin compaction associated with pro-inflammatory polarization is sufficient to increase mechanical resistance to large nuclear deformation, despite a reduction in lamin A/C levels. Our work is expected to motivate deeper investigations into the nuclear mechanobiology of immune cells and prompt further mechanical characterizations of cells with varying chromatin organization or lamin levels.

## Methods

### Bone marrow-derived macrophage (BMDM) isolation and differentiation

Femurs and tibia were isolated from 6-8 week old C57BL/6 mice using previously established protocols^103^. The ends of the bones were cut with scissors, and the marrow was flushed using 10 mL of Dulbecco’s Modified Eagle Medium (DMEM, Thermo Fisher Scientific, 11966025) in a syringe fit with a 27-gauge needle. Bone marrow was resuspended in DMEM and filtered through a 40 µm cell strainer (Greiner Bio-One, 542040). Bone marrow was centrifuged at 400g for 5 minutes to pellet cells, and cells were resuspended at 5 million cells/mL in Recovery Cell Culture Freezing Media (Life Technologies, 12648010). Cells were frozen in liquid nitrogen for a maximum of 4 months. To differentiate the bone marrow cells, 5 million cells were thawed into DMEM and centrifuged to pellet the cells to remove the freezing media. Cells were resuspended in 10 mL of BMDM media containing DMEM, 10% heat inactivated fetal bovine serum (FBS, Avantar, 76509-324), 1% penicillin and streptomycin (Thermo Fisher Scientific, 10378016), and 10 ng/mL of macrophage colony stimulating factor (MCSF, Peprotech, 315-02-50UG) and plated on 10 cm (non-TC treated) petri dishes (VWR, 25384-342). On day 2 of differentiation cells were supplemented with 10 ng/mL of MCSF. On day 4 of differentiation, 5 mL of BMDM media was removed and replaced with fresh BMDM media and supplemented with 10 ng/mL of MCSF. Day 7 differentiated BMDM were used for all experiments unless otherwise stated. BMDM were maintained at 37°C and 5% CO_2_ during differentiation and culture.

### Flow cytometry

F4/80 expression of BMDMs cultured for 6 days was assessed to ensure robust differentiation prior to experiments. BMDM were detached from the petri dishes by incubating the dishes with 5 mL of warm 1x DPBS and gently washing the dish to ensure detachment. Cells were counted and resuspended to a concentration of 2×10^6^ cells/mL in FACS buffer containing 1× Dulbecco’s Phosphate-Buffered Saline (DPBS), 5% heat inactivated FBS, and 2.5 mM EDTA (Thermo Fisher Scientific, AM9260G). 200 µL or 4×10^5^ cells were transferred to Eppendorf tubes to be stained. Samples were stained with 1:000 Live/Dead cell viability kit (Thermo Fisher Scientific, L34973) for 1 hour at 4°C. Samples were washed 3 times with cold FACS buffer. Fc receptors were blocked using FC block rat anti-mouse CD16/CD32 antibody (BD Bioscience 553142) 1:50 for 10 minutes. Cells were stained with F4/80 antibody (Biolegend 123107) or F4/80 isotype control (Biolegend 400505) conjugated to FITC at 1:200 or CD11b antibody (Beckton Dickenson 553397) or CD11b isotype control (Beckton Dickenson 553989) conjugated to PE at 1:100 for 1 hour on ice in the dark. Cells were washed twice with cold FACS buffer and resuspended in a final volume of 500 µL cold FACS buffer. Samples were passed through a 40 µm cell strainer to ensure no cells clumps were present. Flow cytometry was performed using an Accuri C6+ flow cytometer (Becton Dickenson) or a Thermo Fisher Attune NxT analyzer. Samples were analyzed for cell cycle using FlowJo (v10.10). Gating strategies included debris removal and singlet selection. F4/80 and CD11b expression was compared to the isotype control.

### Pro-inflammatory (M1) macrophage polarization

Day 6 differentiated BMDM were detached using warm PBS seeded onto fibronectin coated well plates or #1.5 polymer bottom 18-well coverslips (Ibidi, 81816), and allowed to attach for 2 hours in BMDM media supplemented with 10 ng/mL MCSF. M0 macrophages received 10 ng/mL MCSF alone, while M1 macrophages received BMDM media supplemented with 10 ng/uL MCSF, 100 ng/mL E.coli derived Lipopolysaccharide (LPS, Millipore Sigma, L2630-10MG), and 20 ng/mL interferon-gamma (IFNγ, Peprotech, 315-05-20UG). BMDM were polarized for 24 hours unless otherwise stated prior to experimental use. For all experiments requiring detachment of M1 macrophages (cell cycle analysis, micropipette aspiration, and transwell migration assays), these cells and their respective M0 controls were detached by incubating with accutase for 5 minutes at room temperature.

### DNA Content and Cell Cycle Analysis

D6 differentiated BMDMs were polarized to M1 macrophages or kept as M0 macrophages for 24 hours. BMDMs were detached from the petri dishes by incubating the dishes with 5 mL of accutase at room temperature and gently washing the dish to ensure detachment. Samples were washed with 1× PBS and resuspended in 500 µL PBS. To fix the samples, 4.5 mL of cold 70% ethanol was added to each sample while gently vortexing. The samples were incubated at 4°C for 1 hour and then washed 3 times with 1× PBS. Samples were resuspended in 1 mL in solution containing 3.8 mM sodium citrate, 50 µg/mL propidium iodide (PI, Biotium, #40017), and 10 µg/mL RNase A (ThermoFisher, R1253) in 1× PBS overnight. The next morning, samples were washed 3 times with 1× PBS, resuspended in in FACS buffer containing 1× Dulbecco’s Phosphate-Buffered Saline (DPBS), 5% heat inactivated FBS, and 2.5 mM EDTA (Thermo Fisher Scientific, AM9260G), and passed through a 40 µm cell strainer to ensure that no cells clumps were present. Samples were run on a ThermoFisher Attune NxT analyzer. M0 and M1 macrophage populations were identified using SSC-A vs. FSC-A gating, and singlet cells were gated using FSC-H vs. FSC-A plots (Supplementary Figure 1B, C). Samples were excited at 488 nm to identify PI broad emission centered around 600 nm^104,105^. Samples were analyzed for DNA content and cell cycle distribution using FlowJo (v10.10), and cell cycle phases were deconvolved using the Cell Cycle analysis tool with the Watson (pragmatic) model. G1 and G2 peak widths were equalized and set to minimize the root mean square deviation (RMSD).

### Immunostaining

Day 6 differentiated BMDMs (12×10^3^) were seeded on fibronectin coated 18-well Ibidi coverslips. Cells were either left unpolarized or polarized to M1 BMDM for 24 hours. Cells were fixed with 4% paraformaldehyde in PBS for 15 minutes, followed by three 5-minute washes with PBS. Cells were incubated for 1 hour in IF permeabilization + blocking buffer consisting of 0.1% Triton X-100, 0.05% Tween 20, 5% BSA, 5% donkey or goat serum (species the secondary antibody was raised in) and 1:50 FC Block in PBS. Primary antibodies were diluted in IF blocking buffer containing 5% BSA, 5% donkey or goat serum and 1:50 FC Block in PBS and incubated overnight at 4°C. Primary antibodies used included iNOS (1:100, Cell Signaling, 13120S), lamin A (1:100, Santa Cruz, sc-518013), lamin B1 (1:100, Santa Cruz, sc-374015), Histone-3 (1:100, Santa Cruz, sc-517576), H3k9me3 (1:100, Abcam, ab8898), H3k27me3 (1:100, Millipore Sigma, 07-499). Cells were washed 3 times for 5 minutes each in PBS, and secondary antibody solution was added at 1:250 dilution for 1 hour at room temperature. Secondary antibodies used were Alexa Fluor 488, 568, or 647-conjugated donkey anti mouse or rabbit-IgG antibodies (Invitrogen). Cells were stained with Spy555- Actin (Cytoskeleton, CY-SC202) at 1:1000 dilution in PBS and DAPI (Thermo Fisher Scientific, 62247) at 1:500 dilution in PBS at room temperature for 1 hour. Samples were washed 3 times for 5 minutes each in PBS and left in PBS +0.02% sodium azide in the dark until imaging.

### Protein extraction and immunoblotting

Day 6 differentiated BMDMs (3×10^5^) were seeded on fibronectin coated 6-well plates and allowed to attach. BMDM were either left unpolarized or polarized to M1 BMDMs for 24 hours. 4T1 and mouse embryonic fibroblasts (MEFs) were seeded at 3×10^5^ cells in a 6-well plate for 24 hours. All cells were washed with cold 1× PBS and lysed on ice using high salt RIPA buffer (12 mM sodium deoxycholate, 50 mM Tris-HCl pH 8.0, 750 mM NaCl, 1% (v/v) NP-40 alternative and 0.1% (v/v) SDS in ultrapure water). To extract Lamins, samples were vortexed for 5 minutes at 4°C, sonicated for 30 seconds at 36% amplitude, boiled for 2 min at 95°C, and centrifuged at 4°C for 15 minutes at 14,000×g. The supernatant was stored at –70°C. Protein concentration was determined using a Bradford Assay. Equal amounts (40 µg) of each sample was diluted in 5× Laemmli Buffer, denatured for 3 minutes at 95°C, loaded into a 4-12% Bis-Tris gel (Invitrogen NP0322) and run for 1.5 hours at 100 V. Proteins were transferred onto a PVDF membrane for 1.5 hours at 16 V. Membranes were blocked using 5% BSA in Tris-buffered saline plus 1% Tween-20 (TBST). Primary antibodies were incubated overnight at 4°C in TBST+5% BSA. Primary antibodies include Lamin A/C (1:1000, Santa Cruz, sc-376248), Lamin B1 (1:1000, Santa Cruz, sc-374015), lamin B1 (1:1000 Proteintech, 12987-1-AP), emerin (1:1000, Leica, 50-255-2258), GAPDH (1:1000, Cell Signaling, 21180S), LBR (1:1000, abcam, AB232731), H3 (1:2000 Santa Cruz, sc-517576).

Simultaneously, the gel was stained with 20 mL Coomassie (Bio-Rad 1610786) overnight. Membranes were washed 3×10 minutes using TBST. Secondary antibody Licor IRDye 680RD donkey anti-mouse-IgG (926-68072; 1:5000) was added and incubated for 1 hour at room temperature. Following 3×10 min TBST washes, the membrane was imaged using the Odessey Licor scanner. Brightness and contrast were adjusted using Image Studio Lite (Version 5.2) software. After Coomassie staining, the gel was imaged with a Gel Doc XR+ with ImageLab Software (Bio-Rad). Band intensities and total protein intensities were quantified using FIJI. Quantification is reported as band intensity/ total protein intensity. All full blots and total protein gels are shown in Supplementary Figure 3.

### Micropipette aspiration assay

Micropipette aspiration was performed according to a previously reported protocol^66^. On day 6 of differentiation, ∼5×10^6^ BMDMs were seeded on petri dishes and were left unpolarized as M0 BMDMs or polarized to M1 BMDMs for 6 and 24 hours. After polarization, cells were detached using accutase (VWR, 10210-214) and resuspended in 1× DPBS supplemented with 2% BSA, 0.2% FBS, and 10 mM EDTA to prevent cell clumping. For experiments with cytochalasin D treatment, cells were treated with 4 µM of cytochalasin D or vehicle control (water) for 20 minutes in suspension. Cells were stained with 10 µM/mL Hoechst 33342 prior to micropipette aspiration. During micropipette aspiration, cells flow through a microfluidic device and are trapped in small pockets due to a pressure gradient across the device. Timelapse image acquisition allows for visualization of nuclear deformability into the pockets. Images are acquired every 5 seconds for 40 frames. After image acquisition the cells in the pockets can be cleared for subsequent acquisition of more nuclei. Nuclear protrusion length per frame is calculated by a custom-written MatLab script, available upon request. Data are plotted as nuclear protrusion length over time and average protrusion length at specific time points to perform statistical analysis.

### Transwell migration assay

On day 6 of differentiation, ∼5×10^6^ BMDMs were seeded on petri dishes and were left unpolarized as M0 BMDMs or polarized to M1 BMDMs for 6 and 24 hours.

Cells were detached using accutase and resuspended to a final concentration of 1×10^6^ cells/mL. 50 µL (50,000 cells) of either M0 or M1 cell solution was seeded into the top insert of either 3 µm, 5µm, or 8 µm polycarbonate transwell membranes (Corning, catalog# 3415, 3421, 3422, respectively) and allowed to incubate for 15 minutes at 37°C to allow cells to settle in the transwell insert. After incubation, 700 µL of either M0 or M1 BMDM media was slowly added to the bottom well. For experiments using a chemoattractant, 100 nM N-Formylmethionyl-leucyl-phenylalanine (fMLP) or vehicle control was added to the M0 or M1 media before adding it to the bottom well. After either 6 or 24 hours of migration, cell media was removed and transwell inserts were washed with 1× PBS. Cells were fixed by adding 4% paraformaldehyde (PFA) to the top and bottom of the transwell inserts and incubated at room temperature for 15 minutes. After fixation, PFA was removed and transwell inserts were washed with 1× PBS and stored in PBS at 4°C. To count nuclei, transwell inserts were stained with 1:500 DAPI for 30 minutes and samples were imaged using a Zeiss LSM 900 confocal microscope with the AiryScan module and a 40×/0.8 NA water immersion objective. For each replicate, 6-8 fields of view from two transwell devices were imaged. A CellProfiler (version 4.2.6) pipeline was used to count nuclei. Only cells with nuclei that had completely migrated through the transwell inserts were counted, and any cells with nuclei that had only partially migrated through the pores were manually excluded.

### Fluorescence and confocal microscopy

iNOS staining and micropipette data was acquired using an inverted Zeiss Observer Z1 epifluorescence microscope with Hamamatsu Orca Flash 4.0 camera. Confocal microscopy images used for all immunofluorescence and nuclear volume/morphology analysis were acquired using a Zeiss LSM 900 confocal microscope with the Airyscan module and a 40×/0.8 NA water immersion objective. The optimal z-stack slice determined by Zen Blue (Zeiss) software was used for all 3D imaging.

### Immunofluorescence and nuclear morphology image analysis

Intensity profile line plots were computed using a custom FIJI macro available upon request. This macro plots the fluorescence intensity across a line drawn across a z-slice through the nucleus. The program converts the line length into relative nuclear distances to account for different sized nuclei. 3D nuclear volume and height were analyzed using Arivis Pro software (version 4.2). Nuclei were segmented based on the DAPI signal. The Arivis pipeline is available upon request. All other quantitative fluorescence intensity and nuclear morphology measurements were analyzed using CellProfiler (version 4.2.6) software. The pipeline performs a background subtraction and identifies cells first based on DAPI thresholding and second based on Spy555-Actin staining if included. The mean intensity of each channel was calculated, as well as nuclear area, form factor, and cytoplasmic major and minor axis length. The cytoplasmic aspect ratio was calculated as major axis/minor axis. To calculate the proportions of nuclei with circumferential nuclear envelope wrinkling, nuclear envelope invaginations, and nuclear envelope tunnels, images were blinded and individual nuclei were manually scored, with immunofluorescence labeling for lamin B1 used to identify the nuclear envelope. Nuclear invaginations were identified as indentations in the nuclear envelope using the orthogonal projection of 3D stack images; tunnels were identified as nuclear envelope features that transverses the entire nucleus between the basal and apical surface; and circumferential wrinkling was identified as wrinkling around the periphery of the nucleus using x-y max intensity projections of the 3D z-stack images. The percentage of excess perimeter was calculated by

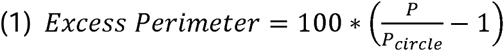

where P is the perimeter of the nucleus calculated using Cell Profiler, and P_CirCle_ is calculated as the perimeter of a perfect circle with the equivalent nuclear area A by,

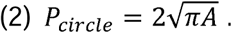

For analysis of nuclear chromocenter size and distribution in M0 and M1 macrophages, 2D maximum intensity projections (x-y plane) of 3D z-slice images of nuclei stained with DAPI were imported into CellProfiler. Chromocenters were thresholded based on the DAPI signal and each chromocenter was assigned to the corresponding ‘parent’ nucleus. Nearest neighbor calculations were produced by the CellProfiler “MeasureObjectNeighbors” module, whereas the intensity distribution of chromocenters was measured using the “MeasureObjectIntensityDistribution” module.

### Brillouin microscopy

Day 6 differentiated BMDMs (2×10^5^) were seeded on fibronectin coated Ibidi coverslips. Cells were either left unpolarized or polarized to M1 BMDM for 24 hours. BMDMs were stained with Hoechst for 30 minutes prior to performing Brillouin Microscopy. A custom built inverted confocal Brillouin microscope was used to acquire the Brillouin frequency shift of M0 and M1 BMDMs. Samples were incubated at 37°C with 5% CO_2_ during imaging using a stage-top Okolab Incubator. Data was acquired using A 660 nM laser (Torus, Laser Quantum Inc.) illuminating the sample with a power of 20 mW. Samples were focused using a 40× objective (Olympus, NA = 0.95). The backscattered Brillouin frequency was collected by the same objective analyzed by two stage virtually imaged phase array-based spectrometer with 15 GHz of free spectral range. The Brillouin spectrum was recorded by an electron charged multiplying charge coupled device camera (iXon Andor) with an exposure time of 0.05 seconds. 2D Brillouin images of cells were acquired using an XY motorized stage with a step size of 1 µm. A brightfield microscope with a CMOS camera (Andor Neo) co-aligned with the Brillouin scanning arm was used to acquire brightfield, fluorescent and nuclear positions using the Hoechst signal.

### Brillouin data acquisition and analysis

LabVIEW-based acquisition program (National Instruments, version 2024) was developed in-house to capture brightfield images, fluorescence images, and Brillouin spectra. Spectrometer was calibrated using electro-optic modulator allowing determination of the free spectral range and the pixel-to-frequency conversion factor. Brillouin shifts at each pixel were extracted by fitting the acquired spectra to a Lorentzian function using MATLAB (MathWorks, R2024a). The resulting pixel-wise Brillouin shift values were used to reconstruct two-dimensional Brillouin images. Co-registration of Brillouin images with corresponding brightfield and fluorescence images was conducted in MATLAB. Brightfield and fluorescence images were resized to match the spatial resolution of the Brillouin images, aligned accordingly, and overlaid to delineate nuclear regions. This co-registration enabled precise extraction and quantification of Brillouin shifts of the cell nucleus. The average Brillouin frequency shift of the nucleus was calculated for all cells.

### Fluorescence lifetime imaging (FLIM)

Day 6 differentiated BMDMs (12×10^3^) were seeded on fibronectin coated 18-well Ibidi coverslips. Cells were either left unpolarized or polarized to M1 BMDM for 24 hours. Prior to imaging cells were stained with 5 µM/mL Hoechst 33342 for 30 minutes and media was changed to Fluorobrite DMEM supplemented with 10% heat inactivated FBS, penicillin and streptomycin (50 U/ml, Thermo Fisher Scientific), GlutaMAX, 10 mM HEPES (Gibco), 10 ng/mL MCSF, and LPS + IFNγ for M1 polarization conditions.

Fluorescence lifetime imaging microscopy (FLIM) was performed using a Zeiss LSM 880 inverted confocal microscope with a 63×/1.4 NA oil immersion objective. Samples were excited at 800 nm using a Spectra-Physics Insight DS femtosecond source (∼120 fs at an 80 MHz repetition rate). Fluorescence was passed through a BGG22 filter (Chroma Technology, Rockingham, VT) and detected using a GaAsP photomultiplier tube (PMT) operating in photon counting mode in the R4 channel of the LSM 880 non-descanned detector. Laser power was adjusted to limit the photon detection rate to ∼0.2% of the repetition rate. FLIM data was acquired using an external PC with a SPC-820 time-correlated single photon counting (TCSPC) module (Becker & Hickl, Berlin) and the SPCImage analysis software (Becker & Hickl, Berlin). Using a 1.54 μs pixel dwell time, photon counts were acquired for 60 seconds per FLIM image (∼60 frames) and binned to 256×256 from 512×512. Data was fit to a single exponential model using the “Incomplete Multiexponential” model provided by the software to account for the 80 MHz repetition rate. Fluorescence lifetime per pixel was calculated in SPCImage and exported to Fiji for further analysis where floating-point images were created from the data. The mean fluorescence lifetime (ns) per nucleus was calculated by thresholding the image based on the Hoechst signal and applying this mask to the float-point lifetime images.

### Chromatin dynamics acquisition and analysis

Day 6 differentiated BMDMs (12×10^3^) were seeded on fibronectin coated 18-well Ibidi coverslips. Cells were either left unpolarized or polarized to M1 BMDMs for 24 hours. Prior to imaging, cells were stained with 5.0 µM/mL Hoechst 33342 for 30 minutes. Time-lapse videos of individual nuclei were acquired for a total of 1 minute imaging every 200 ms using a Zeiss LSM 900 confocal microscope with the AiryScan module and a 63x/1.4 NA oil immersion objective. Time-lapse images were stabilized using Fiji (v1.54p) to correct for both rotational and translational drift. Chromatin puncta (i.e., chromocenters) were identified as single particles (spots) in the stabilized frames via a Laplacian of Gaussian (LoG) detector, configured with a radius of 0.85 µm and a threshold of 50.0. Spot trajectories were generated using the Simple LAP tracker with the following parameters: maximum linking distance of 3 µm, gap-closing distance of 5 µm, and a maximum frame gap of 2. Only tracks containing more than 298 spots, indicating particle presence in at least 298 out of 300 frames, were retained for downstream analysis. All spot identification and tracking was performed using Trackmate (version 7.12.2) as ImageJ/Fiji plugin^106,107^. To calculate the ensemble-time-averaged mean squared displacement (MSD) for each condition, we imported each particle tracks within that condition to either previously developed GEMSpa software^108^ or custom Matlab (R2019a) scripts (https://github.com/liamholtlab/Nuclear_mechanics_analyses) using the following formulas.

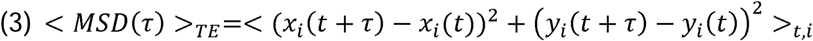

By plotting the log_10_(MSD) versus log_10_(*τ*), we derived local alpha a(r) values from the slope of linear fitting using a sliding window approach (with a sliding window including 5 neighboring points) and calculated over iterations of regions along the plot.

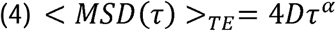

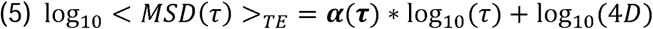

To further characterize the material properties of the chromatin based on chromatin puncta movement, we calculated the elastic and viscoelastic properties through complex modulus using the generalized Stokes-Einstein relationship for viscoelastic materials^109,110^ given by the equations:

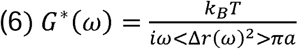

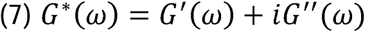

where ω is the frequency domain, *<*ΔJ*r(*ω*)^2^ >* is the Fourier transform of MSD, and a is the tracer particle radius. *G*(*ω) can be separated into the elastic component as the storage modulus *G’(*ω) and the viscous component as the loss modulus *G’’(*ω). To account for ATP-driven non-equilibrium process in chromatin, we used an effective temperature *T* rather than the actual environmental temperature^111^. Since values of *T* and *a* are unknown, we calculated c^’^(w)a/T and c“(w)a/T and compared M1 and M0 cells to evaluate their relative elastic modulus and viscous modulus.

### Statistical Analysis

Statistical analyses were performed using GraphPad Prism (10.6.0). For comparison of 2 groups, a paired *t*-test was performed. For groups of 3, we computed statistical significance using one-way ANOVA with a Dunnett’s test or Tukey’s honestly significant difference (HSD) test for multiple comparisons. A two-way ANOVA was used when comparing the effect of two independent variables. For all quantification of single cell data, data is plotted showing the replicate means in large icons, with individual data points plotted using light gray circles. Horizontal lines indicate the mean ± standard error of the mean (SEM). All data represent three individual replicates from BMDM derived from both male and female mice unless noted otherwise. All quantitative analysis data was blinded using Fiji Blind Analysis Tools plugin when possible.

### Animals

For bone marrow collection, mice were anesthetized using Isoflurane (Covetrus catalog# 11695-6777-2). Immediately following cervical dislocation, the femurs and tibia were collected. All experiments, including breeding, maintenance, and euthanasia of animals, were performed in accordance with relevant guidelines and ethical regulations approved by the Cornell University Institutional Animal Care and Use Committee (IACUC protocol 2011–0099). Mice were housed in a disease-free barrier facility with 12/12 h light/dark cycles.

## Acknowledgements

This work was performed in part at the Cornell NanoScale Facility, a member of the National Nanotechnology Coordinated Infrastructure (NNCI), which is supported by the National Science Foundation (Grant NNCI-2025233). We thank the Biotechnology Resource Center (BRC) Flow Cytometry Facility (RRID: SCR_021740) and sequencing facility (RRID: SCR_021727) at the Cornell Institute of Biotechnology for their resources and technical assistance. This work was supported by awards from the Volkswagen Foundation (A130142 to J.L.), the National Institutes of Health (R01 AR084664, R01 HL082792, R01 GM137605, and R35 GM153257 to J.L., and R01 GM157084 to G.S.), the National Science Foundation (URoL2022048 to J.L.), the Maryland Stem Cell Research Fund (Discovery Grant to G.S.) and the Leducq Foundation (20CVD01 and 24CVD03 to J.L.). The content of this manuscript is solely the responsibility of the authors and does not necessarily represent the official views of the National Institutes of Health. The funding sources played no role in study design, data collection, analysis and interpretation of data, or the writing of this manuscript.

## Data Availability

The datasets generated and/or analyzed during the current study are not publicly available due to data quantity and file size but are available from the corresponding author on reasonable request.

## Code Availability

The code and analysis pipelines used in this study are available by the corresponding author upon reasonable request.

## Author Contributions

This work was conceived by M.E. and J.L. M.E. designed and performed experiments, analysis, and wrote the manuscript. J.O. performed micropipette experiments with cytochalasin D and contributed to analysis, discussion, and interpretation of results. S.H. contributed to chromatin tracking data analysis and flow cytometry. T.S. contributed to chromatin tracking data analysis and complex shear modulus analysis. Y.S.A. performed Brillouin microscopy experiments and analysis. H.S. contributed to Brillouin microscopy experiments. W.R.Z contributed to FLIM experiments, analysis, and data interpretation. G.S. and G.F.W. contributed to Brillion microscopy data interpretation. L.J.H contributed to chromatin tracking experimental design and interpretation. J.L contributed to the experimental design, support and interpretation of analysis, and manuscript revisions. All authors reviewed the manuscript and contributed to the editing.

## References

1. Khatau SB, Hale CM, Stewart-Hutchinson PJ, et al. A perinuclear actin cap regulates nuclear shape. Proc Natl Acad Sci. 2009;106(45):19017–19022. doi:10.1073/pnas.0908686106

2. Lammerding J, Wolf K. Nuclear envelope rupture: Actin fibers are putting the squeeze on the nucleus. J Cell Biol. 2016;215(1):5–8. doi:10.1083/jcb.201609102

3. Versaevel M, Grevesse T, Gabriele S. Spatial coordination between cell and nuclear shape within micropatterned endothelial cells. Nat Commun. 2012;3(1):671. doi:10.1038/ncomms1668

4. Cho S, Vashisth M, Abbas A, et al. Mechanosensing by the Lamina Protects against Nuclear Rupture, DNA Damage, and Cell-Cycle Arrest. Dev Cell. 2019;49(6):920–935.e5. doi:10.1016/j.devcel.2019.04.020

5. Lorber D, Rotkopf R, Volk T. A minimal constraint device for imaging nuclei in live Drosophila contractile larval muscles reveals novel nuclear mechanical dynamics. Lab Chip. 2020;20(12):2100–2112. doi:10.1039/D0LC00214C

6. Friedl P, Weigelin B. Interstitial leukocyte migration and immune function. Nat Immunol. 2008;9(9):960–969. doi:10.1038/ni.f.212

7. Friedl P, Wolf K, Lammerding J. Nuclear mechanics during cell migration. Curr Opin Cell Biol. 2011;23(1):55–64. doi:10.1016/j.ceb.2010.10.015

8. Davidson PM, Denais C, Bakshi MC, Lammerding J. Nuclear deformability constitutes a rate-limiting step during cell migration in 3-D environments. Cell Mol Bioeng. 2014;7(3):293–306. doi:10.1007/s12195-014-0342-y

9. Herbert SP, Stainier DYR. Molecular control of endothelial cell behaviour during blood vessel morphogenesis. Nat Rev Mol Cell Biol. 2011;12(9):551–564. doi:10.1038/nrm3176

10. Weigelin B, Bakker GJ, Friedl P. Intravital third harmonic generation microscopy of collective melanoma cell invasion. IntraVital. 2012;1(1):32–43. doi:10.4161/intv.21223

11. Chepizhko O, Armengol-Collado JM, Alexander S, et al. Confined cell migration along extracellular matrix space in vivo. Proc Natl Acad Sci. 2025;122(1):e2414009121. doi:10.1073/pnas.2414009121

12. Karling T, Weavers H. Immune cells adapt to confined environments in vivo to optimise nuclear plasticity for migration. EMBO Rep. 2025;26(5):1238–1268. doi:10.1038/s44319-025-00381-0

13. Nourshargh S, Alon R. Leukocyte Migration into Inflamed Tissues. Immunity. 2014;41(5):694–707. doi:10.1016/j.immuni.2014.10.008

14. Kalukula Y, Stephens AD, Lammerding J, Gabriele S. Mechanics and functional consequences of nuclear deformations. Nat Rev Mol Cell Biol. 2022;23(9):583–602. doi:10.1038/s41580-022-00480-z

15. Pajerowski JD, Dahl KN, Zhong FL, Sammak PJ, Discher DE. Physical plasticity of the nucleus in stem cell differentiation. Proc Natl Acad Sci. 2007;104(40):15619–15624. doi:10.1073/pnas.0702576104

16. Enyedi B, Jelcic M, Niethammer P. The cell nucleus serves as a mechanotransducer of tissue damage-induced inflammation. Cell. 2016;165(5):1160–1170. doi:10.1016/j.cell.2016.04.016

17. Zwerger M, Jaalouk DE, Lombardi ML, et al. Myopathic lamin mutations impair nuclear stability in cells and tissue and disrupt nucleo-cytoskeletal coupling. Hum Mol Genet. 2013;22(12):2335–2349. doi:10.1093/hmg/ddt079

18. Earle AJ, Kirby TJ, Fedorchak GR, et al. Mutant lamins cause nuclear envelope rupture and DNA damage in skeletal muscle cells. Nat Mater. 2020;19(4):464–473. doi:10.1038/s41563-019-0563-5

19. Venturini V, Pezzano F, Català Castro F, et al. The nucleus measures shape changes for cellular proprioception to control dynamic cell behavior. Science. 2020;370(6514):eaba2644. doi:10.1126/science.aba2644

20. Lomakin AJ, Cattin CJ, Cuvelier D, et al. The nucleus acts as a ruler tailoring cell responses to spatial constraints. Science. 2020;370(6514). doi:10.1126/science.aba2894

21. Long JT, Lammerding J. Nuclear Deformation Lets Cells Gauge Their Physical Confinement. Dev Cell. 2021;56(2):156–158. doi:10.1016/j.devcel.2021.01.002

22. Stephens AD, Banigan EJ, Adam SA, Goldman RD, Marko JF. Chromatin and lamin A determine two different mechanical response regimes of the cell nucleus. Alex R. D, ed. Mol Biol Cell. 2017;28(14):1984–1996. doi:10.1091/mbc.e16-09-0653

23. Furusawa T, Rochman M, Taher L, et al. Chromatin decompaction by the nucleosomal binding protein HMGN5 impairs nuclear sturdiness. Nat Commun. 2015;6(1):6138. doi:10.1038/ncomms7138

24. Aebi U, Cohn J, Buhle L, Gerace L. The nuclear lamina is a meshwork of intermediate-type filaments. Nature. 1986;323(6088):560–564. doi:10.1038/323560a0

25. Wallace M, Zahr H, Perati S, et al. Nuclear damage in LMNA mutant iPSC-derived cardiomyocytes is associated with impaired lamin localization to the nuclear envelope. Mol Biol Cell. 2023;34(12):ar113. doi:10.1091/mbc.E21-10-0527

26. Chen NY, Kim P, Weston TA, et al. Fibroblasts lacking nuclear lamins do not have nuclear blebs or protrusions but nevertheless have frequent nuclear membrane ruptures. Proc Natl Acad Sci. 2018;115(40):10100–10105. doi:10.1073/pnas.1812622115

27. Melcer S, Hezroni H, Rand E, et al. Histone modifications and lamin A regulate chromatin protein dynamics in early embryonic stem cell differentiation. Nat Commun. 2012;3(1):910. doi:10.1038/ncomms1915

28. Bronshtein I, Kepten E, Kanter I, et al. Loss of lamin A function increases chromatin dynamics in the nuclear interior. Nat Commun. 2015;6(1):8044. doi:10.1038/ncomms9044

29. Solovei I, Wang AS, Thanisch K, et al. LBR and Lamin A/C Sequentially Tether Peripheral Heterochromatin and Inversely Regulate Differentiation. Cell. 2013;152(3):584–598. doi:10.1016/j.cell.2013.01.009

30. Mehl JL, Earle A, Lammerding J, Mhlanga M, Vogel V, Jain N. Blockage of lamin-A/C loss diminishes the pro-inflammatory macrophage response. iScience. 2022;25(12):105528. doi:10.1016/j.isci.2022.105528

31. Odell J, Lammerding J. N-terminal tags impair the ability of lamin A to provide structural support to the nucleus. J Cell Sci. 2024;137(16):jcs262207. doi:10.1242/jcs.262207

32. Odell J, Gräf R, Lammerding J. Heterologous expression of Dictyostelium discoideum NE81 in mouse embryo fibroblasts reveals conserved mechanoprotective roles of lamins. Mol Biol Cell. 2024;35(1):ar7. doi:10.1091/mbc.E23-05-0193

33. Swift J, Ivanovska IL, Buxboim A, et al. Nuclear Lamin-A Scales with Tissue Stiffness and Enhances Matrix-Directed Differentiation. Science. 2013;341(6149):1240104. doi:10.1126/science.1240104

34. Guilluy C, Osborne LD, Van Landeghem L, et al. Isolated nuclei adapt to force and reveal a mechanotransduction pathway in the nucleus. Nat Cell Biol. 2014;16(4):376–381. doi:10.1038/ncb2927

35. Lavenus SB, Vosatka KW, Caruso AP, Ullo MF, Khan A, Logue JS. Emerin regulation of nuclear stiffness is required for fast amoeboid migration in confined environments. J Cell Sci. 2022;135(8):jcs259493. doi:10.1242/jcs.259493

36. Chen NY, Kim PH, Tu Y, et al. Increased expression of LAP2β eliminates nuclear membrane ruptures in nuclear lamin–deficient neurons and fibroblasts. Proc Natl Acad Sci. 2021;118(25):e2107770118. doi:10.1073/pnas.2107770118

37. Stephens AD, Liu PZ, Banigan EJ, et al. Chromatin histone modifications and rigidity affect nuclear morphology independent of lamins. Mol Biol Cell. 2018;29(2):220–233. doi:10.1091/mbc.E17-06-0410

38. Dahl KN, Engler AJ, Pajerowski JD, Discher DE. Power-Law Rheology of Isolated Nuclei with Deformation Mapping of Nuclear Substructures. Biophys J. 2005;89(4):2855–2864. doi:10.1529/biophysj.105.062554

39. Manning G, Li A, Eskndir N, Currey M, Stephens AD. Constitutive heterochromatin controls nuclear mechanics, morphology, and integrity through H3K9me3 mediated chromocenter compaction. Nucleus. 2025;16(1):2486816. doi:10.1080/19491034.2025.2486816

40. Nava MM, Miroshnikova YA, Biggs LC, et al. Heterochromatin-Driven Nuclear Softening Protects the Genome against Mechanical Stress-Induced Damage. Cell. 2020;181(4):800–817.e22. doi:10.1016/j.cell.2020.03.052

41. Mosser DM, Edwards JP. Exploring the full spectrum of macrophage activation. Nat Rev Immunol. 2008;8(12):958–969. doi:10.1038/nri2448

42. Sica A, Mantovani A. Macrophage plasticity and polarization: in vivo veritas. March 1, 2012. doi:10.1172/JCI59643

43. Evers TMJ, Sheikhhassani V, Tang H, Haks MC, Ottenhoff THM, Mashaghi A. Single-Cell Mechanical Characterization of Human Macrophages. Adv NanoBiomed Res. 2022;2(7):2100133. doi:10.1002/anbr.202100133

44. Jiao S, Li C, Guo F, et al. SUN1/2 controls macrophage polarization via modulating nuclear size and stiffness. Nat Commun. 2023;14(1). doi:10.1038/s41467-023-42187-5

45. Dooling LJ, Anlaş AA, Tobin MP, et al. Clustered macrophages cooperate to eliminate tumors via coordinated intrudopodia. Proc Natl Acad Sci U S A. 122(27):e2425452122. doi:10.1073/pnas.2425452122

46. Shin SJ, Bayarkhangai B, Tsogtbaatar K, et al. Matrix-Rigidity Cooperates With Biochemical Cues in M2 Macrophage Activation Through Increased Nuclear Deformation and Chromatin Accessibility. Adv Sci. 2025;12(8):2403409. doi:10.1002/advs.202403409

47. Murray PJ, Wynn TA. Obstacles and opportunities for understanding macrophage polarization. J Leukoc Biol. 2011;89(4):557–563. doi:10.1189/jlb.0710409

48. Morris L, Graham CF, Gordon S. Macrophages in haemopoietic and other tissues of the developing mouse detected by the monoclonal antibody F4/80. Development. 1991;112(2):517–526. doi:10.1242/dev.112.2.517

49. Austyn JM, Gordon S. F4/80, a monoclonal antibody directed specifically against the mouse macrophage. Eur J Immunol. 1981;11(10):805–815. doi:10.1002/eji.1830111013

50. Scafidi A, Lind-Holm Mogensen F, Michelucci A. Protocol for the generation and assessment of functional macrophages from mouse bone marrow cells. STAR Protoc. 2025;6(2):103706. doi:10.1016/j.xpro.2025.103706

51. Protocol for the generation and assessment of functional macrophages from mouse bone marrow cells. Accessed February 3, 2026. https://star-protocols.cell.com/protocols/4095

52. McWhorter FY, Wang T, Nguyen P, Chung T, Liu WF. Modulation of macrophage phenotype by cell shape. Proc Natl Acad Sci. 2013;110(43):17253–17258. doi:10.1073/pnas.1308887110

53. Waldo SW, Li Y, Buono C, et al. Heterogeneity of Human Macrophages in Culture and in Atherosclerotic Plaques. Am J Pathol. 2008;172(4):1112–1126. doi:10.2353/ajpath.2008.070513

54. Gravemann S, Schnipper N, Meyer H, et al. Dosage effect of zero to three functional LBR-genes in vivo and in vitro. Nucleus. 2010;1(2):179–189. doi:10.4161/nucl.1.2.11113

55. Vahabikashi A, Adam SA, Medalia O, Goldman RD. Nuclear lamins: Structure and function in mechanobiology. APL Bioeng. 2022;6(1):011503. doi:10.1063/5.0082656

56. Buchwalter A, Kaneshiro JM, Hetzer MW. Coaching from the sidelines: the nuclear periphery in genome regulation. Nat Rev Genet. 2019;20(1):39–50. doi:10.1038/s41576-018-0063-5

57. Marin HC, Allen C, Simental E, et al. The nuclear periphery confers repression on H3K9me2-marked genes and transposons to shape cell fate. Nat Cell Biol. 2025;27(8):1311–1326. doi:10.1038/s41556-025-01703-z

58. Rowat AC, Lammerding J, Ipsen JH. Mechanical Properties of the Cell Nucleus and the Effect of Emerin Deficiency. Biophys J. 2006;91(12):4649–4664. doi:10.1529/biophysj.106.086454

59. Lammerding J, Hsiao J, Schulze PC, Kozlov S, Stewart CL, Lee RT. Abnormal nuclear shape and impaired mechanotransduction in emerin-deficient cells. J Cell Biol. 2005;170(5):781–791. doi:10.1083/jcb.200502148

60. Bell ES, Shah P, Zuela-Sopilniak N, et al. Low lamin A levels enhance confined cell migration and metastatic capacity in breast cancer. Oncogene. 2022;41(36):4211–4230. doi:10.1038/s41388-022-02420-9

61. Odell J, Tang Y, Ambekar YS, et al. The Deformability of the Mammalian Cell Nucleus is Determined by the Identity of the Lamin Rod Domain. bioRxiv. Published online July 22, 2025:2025.07.22.666133. doi:10.1101/2025.07.22.666133

62. Daniel B, Belk JA, Meier SL, et al. Macrophage inflammatory and regenerative response periodicity is programmed by cell cycle and chromatin state. Mol Cell. 2023;83(1):121–138.e7. doi:10.1016/j.molcel.2022.11.017

63. Liu L, Lu Y, Martinez J, et al. Proinflammatory signal suppresses proliferation and shifts macrophage metabolism from Myc-dependent to HIF1α-dependent. Proc Natl Acad Sci. 2016;113(6):1564–1569. doi:10.1073/pnas.1518000113

64. Brändle FB, Frühbauer B, Ceppi I, et al. Nuclear mechanostability emerges from satellite DNA condensation into chromocenters. bioRxiv. Preprint posted online August 8, 2025:2025.08.07.669059. doi:10.1101/2025.08.07.669059

65. Manning G, Li A, Eskndir N, Currey M, Stephens AD. Constitutive heterochromatin controls nuclear mechanics, morphology, and integrity through H3K9me3 mediated chromocenter compaction. Nucleus. 16(1):2486816. doi:10.1080/19491034.2025.2486816

66. Davidson PM, Fedorchak GR, Mondésert-Deveraux S, et al. High-throughput microfluidic micropipette aspiration device to probe time-scale dependent nuclear mechanics in intact cells. Lab Chip. 2019;19(21):3652–3663. doi:10.1039/c9lc00444k

67. Pergola C, Schubert K, Pace S, et al. Modulation of actin dynamics as potential macrophage subtype-targeting anti-tumour strategy. Sci Rep. 2017;7(1):41434. doi:10.1038/srep41434

68. Kabakova I, Zhang J, Xiang Y, et al. Brillouin microscopy. Nat Rev Methods Primer. 2024;4(1):8. doi:10.1038/s43586-023-00286-z

69. Scarcelli G, Yun SH. Confocal Brillouin microscopy for three-dimensional mechanical imaging. Nat Photonics. 2008;2(1):39–43. doi:10.1038/nphoton.2007.250

70. Nikolić M, Scarcelli G. Long-term Brillouin imaging of live cells with reduced absorption-mediated damage at 660 nm wavelength. Biomed Opt Express. 2019;10(4):1567–1580. doi:10.1364/BOE.10.001567

71. Dixon JR, Jung I, Selvaraj S, et al. Chromatin architecture reorganization during stem cell differentiation. Nature. 2015;518(7539):331–336. doi:10.1038/nature14222

72. May D, Yun S, Gonzalez DG, et al. Live imaging reveals chromatin compaction transitions and dynamic transcriptional bursting during stem cell differentiation in vivo. Lakadamyali M, Stainier DY, eds. eLife. 2023;12:e83444. doi:10.7554/eLife.83444

73. Yadav T, Quivy JP, Almouzni G. Chromatin plasticity: A versatile landscape that underlies cell fate and identity. Science. 2018;361(6409):1332–1336. doi:10.1126/science.aat8950

74. Ivashkiv LB. Epigenetic regulation of macrophage polarization and function. Trends Immunol. 2013;34(5):216–223. doi:10.1016/j.it.2012.11.001

75. Zhu Y, van Essen D, Saccani S. Cell-Type-Specific Control of Enhancer Activity by H3K9 Trimethylation. Mol Cell. 2012;46(4):408–423. doi:10.1016/j.molcel.2012.05.011

76. Hachiya R, Shiihashi T, Shirakawa I, et al. The H3K9 methyltransferase Setdb1 regulates TLR4-mediated inflammatory responses in macrophages. Sci Rep. 2016;6(1):28845. doi:10.1038/srep28845

77. Spagnol ST, Dahl KN. Spatially Resolved Quantification of Chromatin Condensation through Differential Local Rheology in Cell Nuclei Fluorescence Lifetime Imaging. PLOS ONE. 2016;11(1):e0146244. doi:10.1371/journal.pone.0146244

78. Booth-Gauthier EA, Alcoser TA, Yang G, Dahl KN. Force-Induced Changes in Subnuclear Movement and Rheology. Biophys J. 2012;103(12):2423–2431. doi:10.1016/j.bpj.2012.10.039

79. Sabri A, Xu X, Krapf D, Weiss M. Elucidating the Origin of Heterogeneous Anomalous Diffusion in the Cytoplasm of Mammalian Cells. Phys Rev Lett. 2020;125(5):058101. doi:10.1103/PhysRevLett.125.058101

80. Paterson N, Lämmermann T. Macrophage network dynamics depend on haptokinesis for optimal local surveillance. Dustin ML, Rothlin CV, Dustin ML, eds. eLife. 2022;11:e75354. doi:10.7554/eLife.75354

81. Vereyken EJ, Heijnen PD, Baron W, de Vries EH, Dijkstra CD, Teunissen CE. Classically and alternatively activated bone marrow derived macrophages differ in cytoskeletal functions and migration towards specific CNS cell types. J Neuroinflammation. 2011;8:58. doi:10.1186/1742-2094-8-58

82. Cui K, Ardell CL, Podolnikova NP, Yakubenko VP. Distinct Migratory Properties of M1, M2, and Resident Macrophages Are Regulated by αDβ2 and αMβ2 Integrin-Mediated Adhesion. Front Immunol. 2018;9:2650. doi:10.3389/fimmu.2018.02650

83. Vogel DY, Heijnen PD, Breur M, et al. Macrophages migrate in an activation-dependent manner to chemokines involved in neuroinflammation. J Neuroinflammation. 2014;11:23. doi:10.1186/1742-2094-11-23

84. Hind LE, Lurier EB, Dembo M, Spiller KL, Hammer DA. Effect of M1–M2 Polarization on the Motility and Traction Stresses of Primary Human Macrophages. Cell Mol Bioeng. 2016;9(3):455–465. doi:10.1007/s12195-016-0435-x

85. Piedra-Quintero ZL, Serrano C, Villegas-Sepúlveda N, et al. Myosin 1F Regulates M1-Polarization by Stimulating Intercellular Adhesion in Macrophages. Front Immunol. 2019;9. doi:10.3389/fimmu.2018.03118

86. Wiesolek HL, Bui TM, Lee JJ, et al. Intercellular Adhesion Molecule 1 Functions as an Efferocytosis Receptor in Inflammatory Macrophages. Am J Pathol. 2020;190(4):874–885. doi:10.1016/j.ajpath.2019.12.006

87. Alonso-Matilla R, Provenzano PP, Odde DJ. Physical principles and mechanisms of cell migration. Npj Biol Phys Mech. 2025;2(1):2. doi:10.1038/s44341-024-00008-w

88. Chen S, Saeed AFUH, Liu Q, et al. Macrophages in immunoregulation and therapeutics. Signal Transduct Target Ther. 2023;8(1):207. doi:10.1038/s41392-023-01452-1

89. Scarcelli G, Polacheck WJ, Nia HT, et al. Noncontact three-dimensional mapping of intracellular hydromechanical properties by Brillouin microscopy. Nat Methods. 2015;12(12):1132–1134. doi:10.1038/nmeth.3616

90. Zhang J, Nou XA, Kim H, Scarcelli G. Brillouin flow cytometry for label-free mechanical phenotyping of the nucleus. Lab Chip. 2017;17(4):663–670. doi:10.1039/C6LC01443G

91. Antonacci G, Braakman S. Biomechanics of subcellular structures by non-invasive Brillouin microscopy. Sci Rep. 2016;6(1):37217. doi:10.1038/srep37217

92. Oishi Y, Manabe I. Macrophages in age-related chronic inflammatory diseases. Npj Aging Mech Dis. 2016;2(1):16018. doi:10.1038/npjamd.2016.18

93. Savage P. Macrophage modulation of tumor immunity. Science. 2024;386(6724):850–851. doi:10.1126/science.adt5661

94. Chen R, Zhang H, Tang B, et al. Macrophages in cardiovascular diseases: molecular mechanisms and therapeutic targets. Signal Transduct Target Ther. 2024;9(1):130. doi:10.1038/s41392-024-01840-1

95. Louwe PA, Badiola Gomez L, Webster H, et al. Recruited macrophages that colonize the post-inflammatory peritoneal niche convert into functionally divergent resident cells. Nat Commun. 2021;12(1):1770. doi:10.1038/s41467-021-21778-0

96. Watanabe S, Alexander M, Misharin AV, Budinger GRS. The role of macrophages in the resolution of inflammation. J Clin Invest. 129(7):2619–2628. doi:10.1172/JCI124615

97. Vinogradov S, Warren G, Wei X. Macrophages associated with tumors as potential targets and therapeutic intermediates. Nanomed. 2014;9(5):695–707. doi:10.2217/nnm.14.13

98. Linke JA, Munn LL, Jain RK. Compressive stresses in cancer: characterization and implications for tumour progression and treatment. Nat Rev Cancer. 2024;24(11):768–791. doi:10.1038/s41568-024-00745-z

99. Moradian H, Roch T, Lendlein A, Gossen M. mRNA Transfection-Induced Activation of Primary Human Monocytes and Macrophages: Dependence on Carrier System and Nucleotide Modification. Sci Rep. 2020;10(1):4181. doi:10.1038/s41598-020-60506-4

100. Khantakova JN, Silkov AN, Tereshchenko VP, Gavrilova EV, Maksyutov RA, Sennikov SV. Transfection of bone marrow derived cells with immunoregulatory proteins. Cytokine. 2018;108:82–88. doi:10.1016/j.cyto.2018.03.028

101. Carralot JP, Kim TK, Lenseigne B, et al. Automated High-Throughput siRNA Transfection in Raw 264.7 Macrophages: A Case Study for Optimization Procedure. SLAS Discov. 2009;14(2):151–160. doi:10.1177/1087057108328762

102. Cheung ST, Shakibakho S, So EY, Mui ALF. Transfecting RAW264.7 Cells with a Luciferase Reporter Gene. JoVE J Vis Exp. 2015;(100):e52807. doi:10.3791/52807

103. Springer NL, Iyengar NM, Bareja R, et al. Obesity-Associated Extracellular Matrix Remodeling Promotes a Macrophage Phenotype Similar to Tumor-Associated Macrophages. Am J Pathol. 2019;189(10):2019–2035. doi:10.1016/j.ajpath.2019.06.005

104. Crissman HA, Steinkamp JA. RAPID, SIMULTANEOUS MEASUREMENT OF DNA, PROTEIN, AND CELL VOLUME IN SINGLE CELLS FROM LARGE MAMMALIAN CELL POPULATIONS. J Cell Biol. 1973;59(3):766–771. doi:10.1083/jcb.59.3.766

105. Krishan A. Rapid flow cytofluorometric analysis of mammalian cell cycle by propidium iodide staining. J Cell Biol. 1975;66(1):188–193. doi:10.1083/jcb.66.1.188

106. Tinevez JY, Perry N, Schindelin J, et al. TrackMate: An open and extensible platform for single-particle tracking. Methods. 2017;115:80–90. doi:10.1016/j.ymeth.2016.09.016

107. TrackMate 7: integrating state-of-the-art segmentation algorithms into tracking pipelines | Nature Methods. Accessed October 12, 2025. https://www.nature.com/articles/s41592-022-01507-1

108. Keegan S, Fenyö D, Holt LJ. GEMspa: a Napari plugin for analysis of single particle tracking data. Biophysics. Preprint posted online June 28, 2023. doi:10.1101/2023.06.26.546612

109. Optical Measurements of Frequency-Dependent Linear Viscoelastic Moduli of Complex Fluids | Phys. Rev. Lett. Accessed October 12, 2025. https://journals.aps.org/prl/abstract/10.1103/PhysRevLett.74.1250

110. Mason TG. Estimating the viscoelastic moduli of complex fluids using the generalized Stokes–Einstein equation. Rheol Acta. 2000;39(4):371–378. doi:10.1007/s003970000094

111. Caragine CM, Kanellakopoulos N, Zidovska A. Mechanical stress affects dynamics and rheology of the human genome. Soft Matter. 2022;18(1):107–116. doi:10.1039/D1SM00983D

