## Supplementary Materials for "Polarization Increases Nuclear Stiffness in Macrophages Despite Reduction in Lamin A/C Levels"

**Supplementary Figures:**

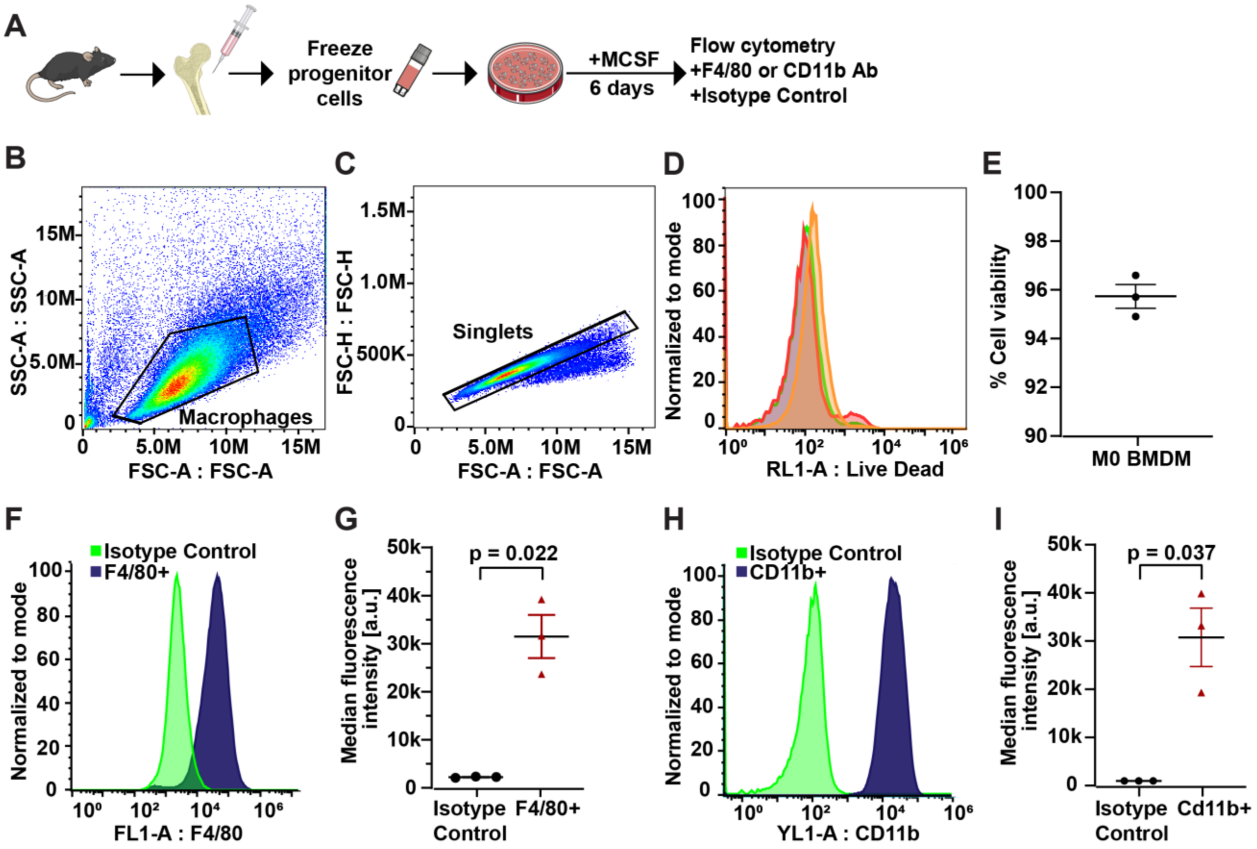


**Supplementary Figure 1. Quantification of F4/80 and CD11b expression by flow cytometry to assess macrophage differentiation.** (**A**) Schematic of macrophage isolation and differentiation timeline. (**B**) Flow cytometry-gating strategy to identify macrophage population using SSC-A vs FSC-A plot. Gated macrophage population outlined in black. (**C**) Flow cytometry-gating strategy to identify macrophage singlets using FSC-H vs FCS-A plot. Gated singlet population outlined in black. (**D**) Histogram of Live/Dead fluorescence profile of 3 replicates of differentiated M0 macrophages. (**E**) Quantification of the percentage of viable macrophages from 3 independent replicates. (**F**) Histogram of F4/80 fluorescence profile of differentiated macrophage stained with F4/80 antibody (navy) compared to isotype control (green). (**G**) Quantification of median F4/80 fluorescence intensity. (**H**) Histogram of CD11b fluorescence profile of differentiated macrophage stained with CD11b antibody (navy) compared to isotype control (green). (**I**) Quantification of median CD11b fluorescence intensity. Data is plotted as a horizontal line at the mean ± SEM, with the replicate means plotted. Statistical analyses in panels G, I are based on paired *t*-tests using replicate means from 3 independent experiments. Panel A icons provided by the Bioicons community from Servier (https://smart.servier.com/) is licensed under CC-BY 3.0.

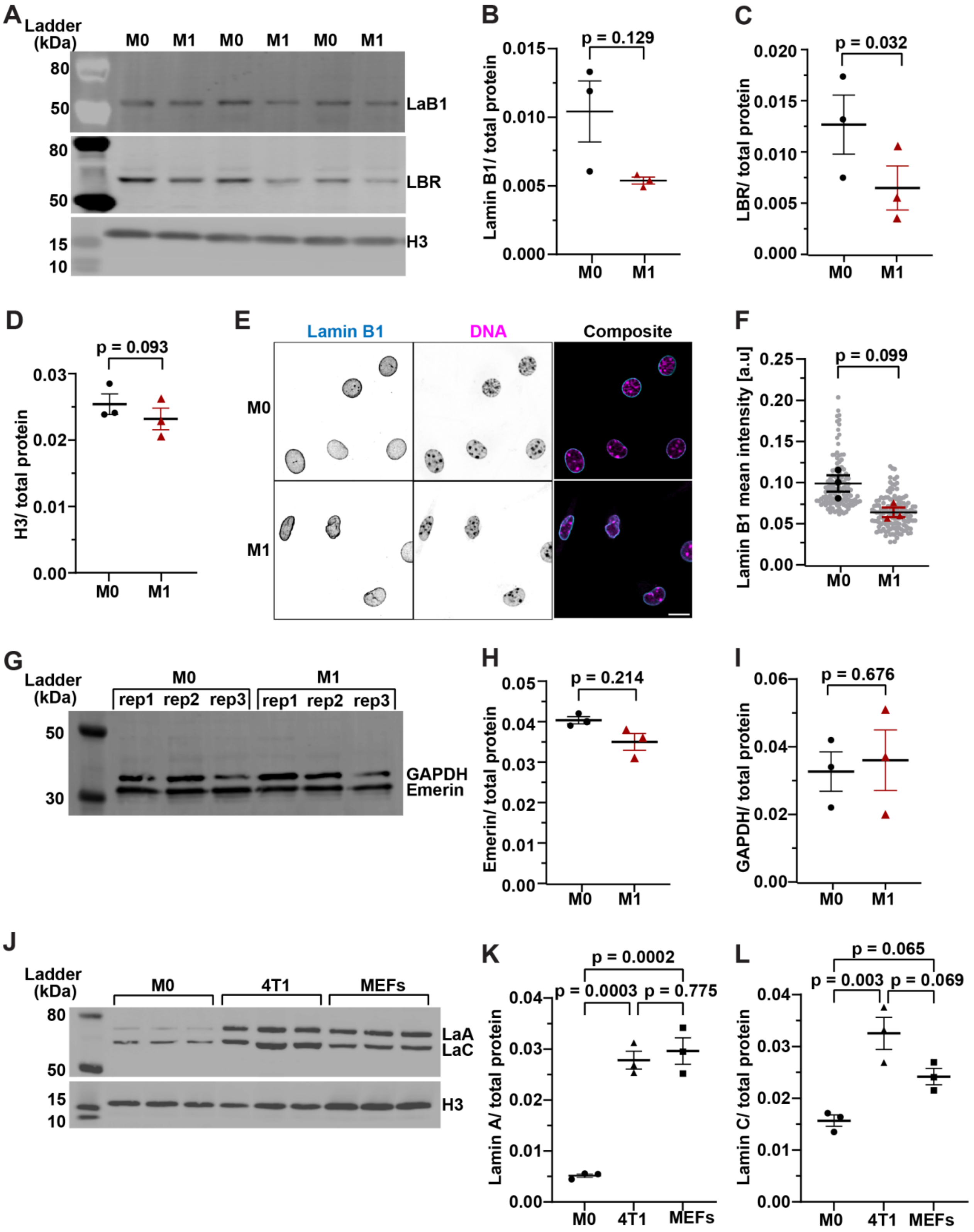

A
Ladder (kDa)
M0 M1 M0 M1 M0 M1
80
50
LaB1
80
50
LBR
15
10
H3
B
p = 0.129
Lamin B1/ total protein
0.015
0.010
0.005
0.000
M0 M1
C
p = 0.032
LBR/ total protein
0.020
0.015
0.010
0.005
0.000
M0 M1
D
p = 0.093
H3/ total protein
0.03
0.02
0.01
0.00
M0 M1
E
Lamin B1
DNA
Composite
M0
M1
F
p = 0.099
Lamin B1 mean intensity [a.u]
0.25
0.20
0.15
0.10
0.05
0.00
M0 M1
G
Ladder (kDa)
M0
M1
rep1 rep2 rep3
rep1 rep2 rep3
50
30
GAPDH
Emerin
H
p = 0.214
Emerin/ total protein
0.05
0.04
0.03
0.02
0.01
0.00
M0 M1
I
p = 0.676
GAPDH/ total protein
0.06
0.04
0.02
0.00
M0 M1
J
Ladder (kDa)
M0
4T1
MEFs
80
50
LaA
LaC
15
10
H3
K
p = 0.0002
p = 0.0003
p = 0.775
Lamin A/ total protein
0.04
0.03
0.02
0.01
0.00
M0 4T1 MEFs
L
p = 0.065
p = 0.003
p = 0.069
Lamin C/ total protein
0.04
0.03
0.02
0.01
0.00
M0 4T1 MEFs

**Supplementary Figure 2. Quantitative analysis of Lamin B1, Lamin B receptor (LBR), Histone-3 (H3), and relative Lamin A/C levels based on immunofluorescence labeling and immunoblot analysis.** (**A**) Representative immunoblots for Lamin B1, LBR, and histone H3 of M0 and M1 macrophages. (**B**) Corresponding quantification of Lamin B1 levels in the immunoblot shown in panel A, normalized to total protein loading. (**C**) Corresponding quantification of LBR levels in the immunoblot shown in panel A, normalized to total protein loading. (**D**) Corresponding quantification of H3 levels in the immunoblots shown in panel A, normalized to total protein loading. (**E**) Representative maximum intensity projection images of cells immunofluorescently labeled for lamin B1. Scale bar = 10 µm. (**F**) Quantification of immunofluorescence images of cells stained for lamin B1. $n$ = 119-135 cells per condition. (**G**) Representative immunoblots for emerin, GAPDH of M0 and M1 macrophages. (**H**) Corresponding quantification of emerin levels in the immunoblot shown in panel G, normalized to total protein loading. (**I**) Corresponding quantification of GAPDH levels in the immunoblot shown in panel G, normalized to total protein loading. (**J**) Representative immunoblots for lamins A/C, H3 of M0 macrophages, 4T1 cells, and MEFs. (**K**) Corresponding quantification of lamin A and levels in the immunoblot shown in panel J, normalized to total protein loading. (**L**) Corresponding quantification of lamin C levels in the immunoblot shown in panel J, normalized to total protein loading. All total protein loading gels are shown in Supplementary Figure 3. For quantification involving individual cell measurements (F), data are shown as horizontal lines for the mean ± SEM, with replicate averages plotted in large black circles (M0) or red triangles (M1), and individual measurements shown as small grey circles. Statistical analyses in panels B-D, F, H, I are based on paired $t$-tests using replicate means from 3 independent experiments.

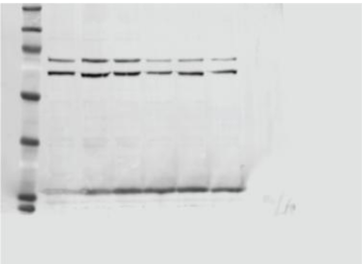

**Fig. 1F full blot (mouse)**
**Antibodies used:**
**Lamin A/C (Santa Cruz sc-376248)**
**H3 (Santa Cruz sc-517576)**

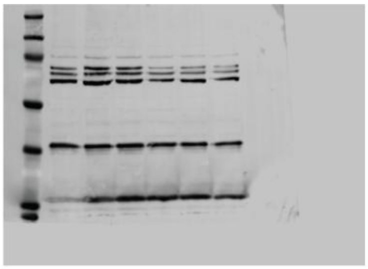

**Supplemental Fig. 2G full blot (mouse)**
**Antibodies used:**
**Lamin A/C (Santa Cruz sc-376248)**
**Lamin B1 (Santa Cruz sc-374015)**
**Emerin (Leica 50-255-2258)**
**H3 (Santa Cruz sc-517576)**

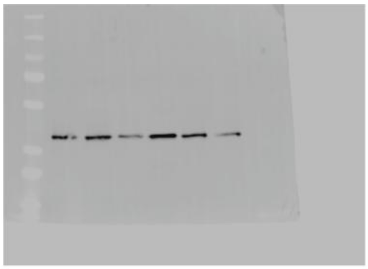

**Supplemental Fig. 2G full blot (rabbit)**
**Antibodies used:**
**GAPDH (Cell Signaling 21180S)**

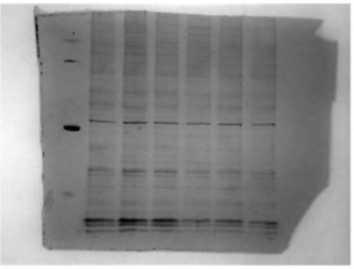

**Fig. 1F, Supplemental Fig 2G**
**total protein**
**coomassie staining**

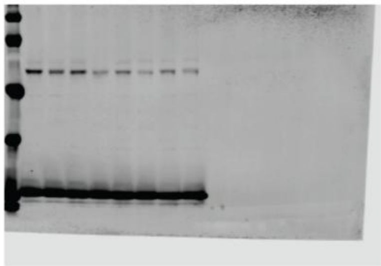

**Supplemental Fig. 2A full blot**
**(mouse, long exposure)**
**Antibodies used:**
**LBR (abcam AB232731)**
**H3 (Santa Cruz sc-517576)**

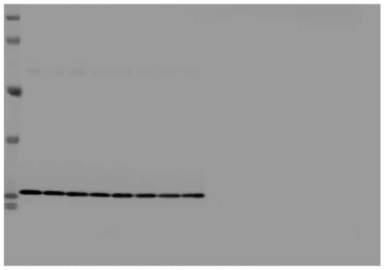

**Supplemental Fig. 2A full blot**
**(mouse, short exposure)**
**Antibodies used:**
**LBR (abcam AB232731)**
**H3 (Santa Cruz sc-517576)**

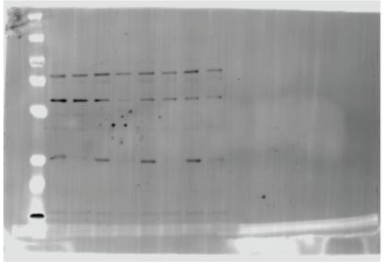

**Supplemental Fig. 2A full blot (rabbit)**
**Antibodies used:**
**Lamin B1 (Proteintech 12987-1-AP)**

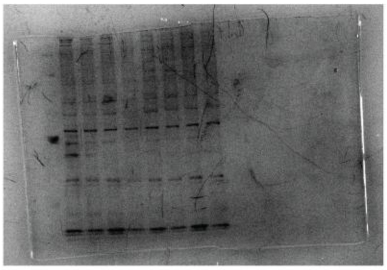

**Supplemental Fig. 2A total protein**
**coomassie staining**

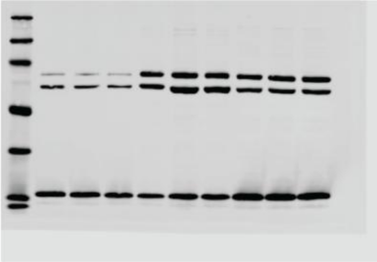

**Supplemental Fig. 2J full blot**
**(long exposure)**
**Antibodies used:**
**Lamin A/C (Santa Cruz sc-376248)**
**H3 (Santa Cruz sc-517576)**

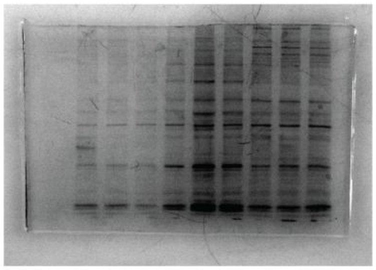

**Supplemental Fig. 2J total protein**
**coomassie staining**

**Supplementary Figure 3. Immunoblot Transparency.** Full, uncropped blots and total protein staining are shown below for each of the immunoblots and gels stained with Coomassie. ThermoFisher Page Ruler plus prestained protein ladder (26617) was used for the molecular weight marker and is present in the first lane of each blot.

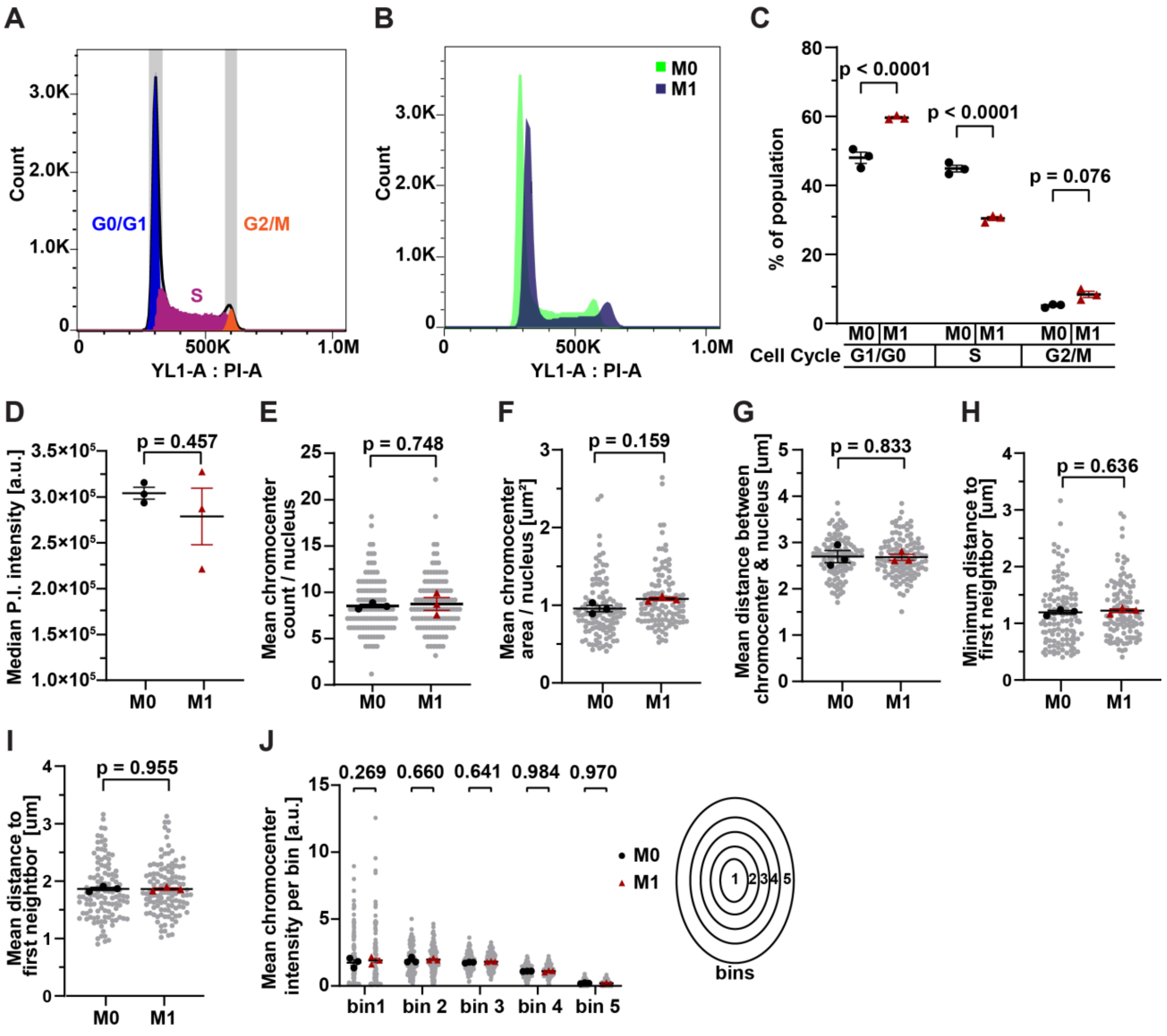


**Supplementary Figure 4. Effect of macrophage polarization on cell cycle and chromocenter distribution.** (**A**) Representative histogram of DNA content from flow cytometry analysis of M0 macrophages based on propidium iodide staining. Cell cycle deconvolution based on DNA content is shown by highlighting the assigned G0/G1 (blue), S (magenta), and G2/M phases (orange). (**B**) Representative histogram of DNA content from flow cytometry analysis of M0 (green) and M1 (blue) macrophages stained with propidium iodide. (**C**) Quantification of cell cycle analysis for M0 and M1 macrophages. (**D**) Quantification of median fluorescence intensity of M0 and m1 macrophages stained with propidium iodide. (**E-G**) Quantification of the mean chromocenters count per nucleus (**E**), total chromocenter area per nucleus (**F**), and mean distance between the chromocenters centroid and nuclear centroid per nucleus in M0 and M1 macrophages (**G**). (**H-I**) Quantification of the minimum (**H**) and mean distance (**I**) to the first neighbor for each centrosome per nucleus. (**J**) Quantification of chromocenter distribution displayed as mean chromocenters intensity per radial nuclear bin, normalized per nuclear area. For all graphs, data is represented as horizontal lines at the mean ± SEM, with replicate averages plotted in large black circles (M0) or red triangles (M1), and individual measurements shown as small grey circles, where applicable.

A two-way ANOVA was performed on the replicate means with Tukey's multiple comparison test to compare means of each group in C, J. A student's *t*-test was performed comparing replicate means from 3 independent experiments in D-I.

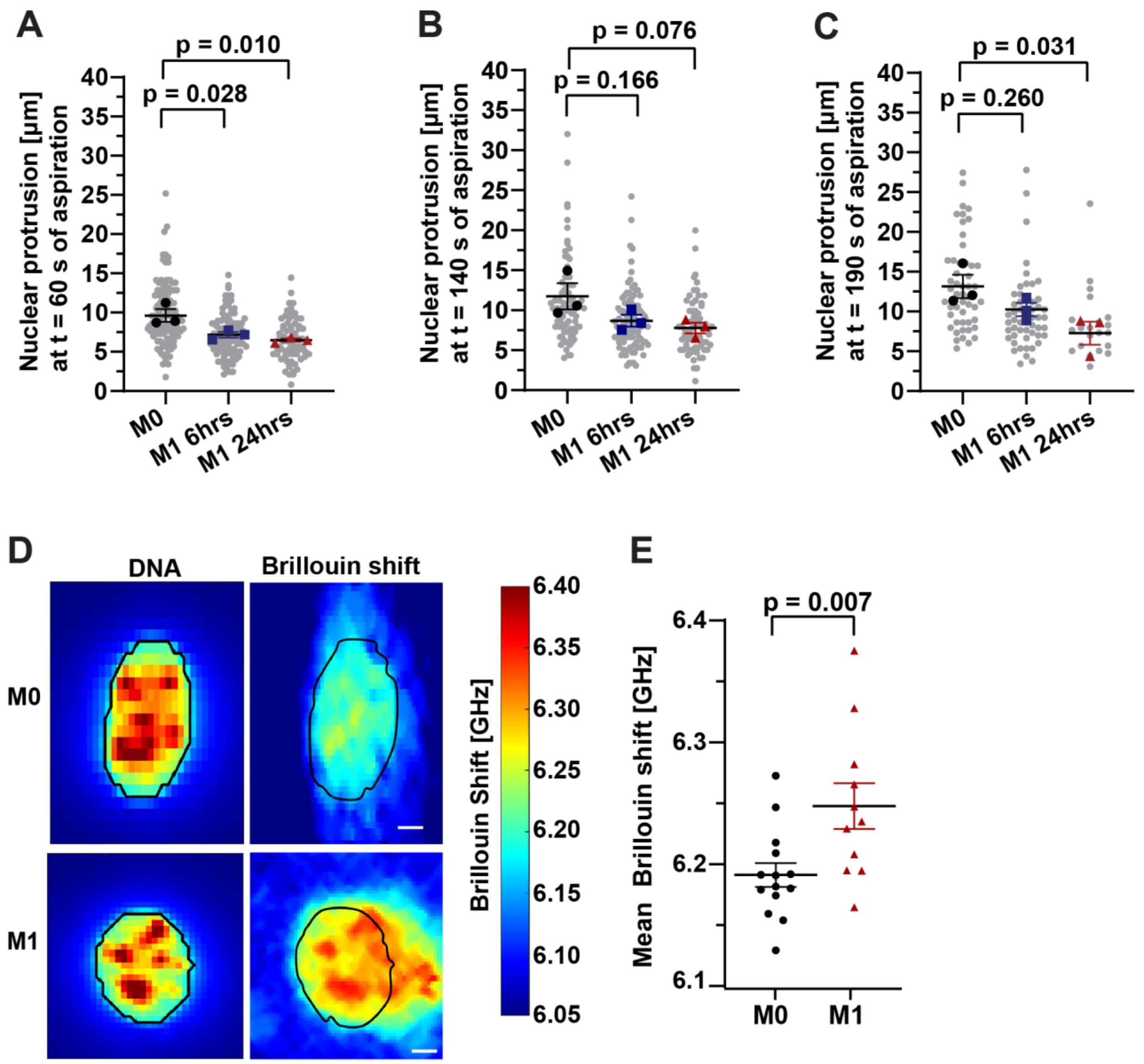


**Supplementary Figure 5. Additional micropipette aspiration time points and Brillouin Microscopy**. Nuclear protrusion length measured at 60 seconds (**A**), 140 seconds (**B**), and 190 seconds (**C**) after the start of nuclear aspiration. Data shown as mean ± SEM, with replicate averages plotted in large black circles (M0), blue squared (M1 6 hrs), or red triangles (M1 24 hrs), and individual measurements shown as small grey circles. (**D**) Representative images showing Hoechst fluorescence intensity (for nuclear segmentation) and Brillouin frequency shift (for mechanical measurements) of M1 and M0 macrophages. (**E**) Quantification of Brillouin frequency shift of M0 and M1 nuclei. Points represent individual cell measurements, and the bars indicate the mean. For panels A-C, statistical analyses are based on one-way ANOVA on the replicate means with Dunnett's multiple comparison test to compare means of each group in A-C. Statistical analysis in panel E is based on student's *t*-tests comparing individual cell measurements. Scale bar = 2 µm.

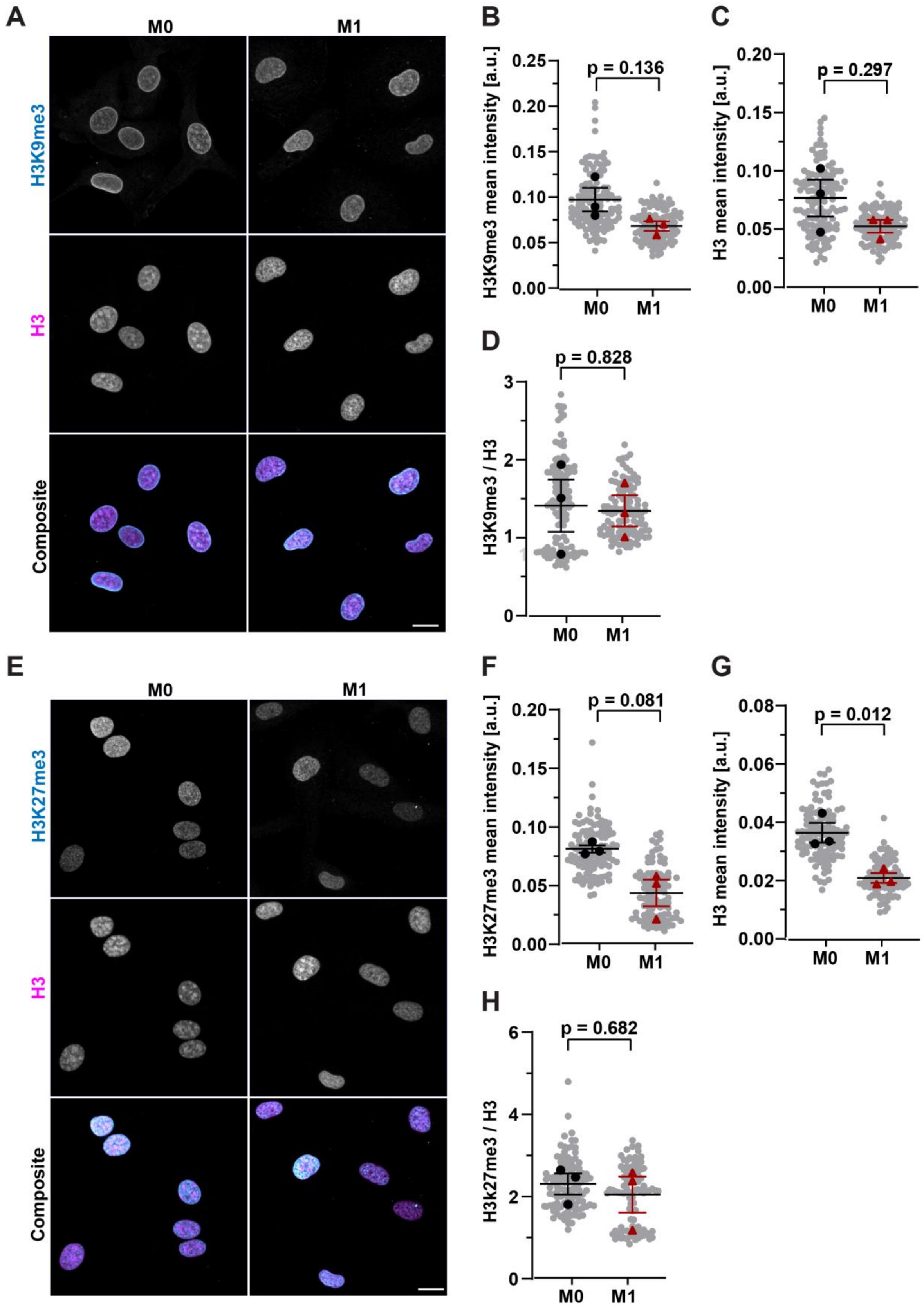
A
M0
M1
H3K9me3
H3
Composite
B
p = 0.136
H3K9me3 mean intensity [a.u.]
M0
M1
C
p = 0.297
H3 mean intensity [a.u.]
D
p = 0.828
H3K9me3 / H3
E
H3K27me3
F
p = 0.081
H3K27me3 mean intensity [a.u.]
G
p = 0.012
H
p = 0.682
H3k27me3 / H3

**Supplementary Figure 6. Global heterochromatin levels do not change significantly in response to macrophage polarization.** (**A**) Representative maximum intensity projection images of cells immunofluorescently labeled for trimethylated histone H3K9 (H3k9me3) and total histone H3. Scale bar = 10 μm (**B-D**) Quantification of immunofluorescence images of cells stained for H3k9me3 (**B**), H3 (**C**), and the ratio of H3k9me3 to total H3 (**D**). $n$ = 94-117 cells per condition from three independent experiments. (**E**) Representative maximum intensity projection images of cells immunofluorescently labeled for trimethylated H3K27 (H3k27me3) and total histone H3. Scale bar = 10 μm (**F-H**) Quantification of immunofluorescence images of cells stained for H3k27me3 (**F**), H3 (**G**), and the ratio of H3k27me3 to total H3 (**H**). $n$ = 76-93 cells per condition. Quantification is represented as a line at the mean ± SEM, with replicate averages plotted in large black circles (M0) or red triangles (M1), and individual measurements shown as small grey circles. Statistical analyses are based on paired $t$-test using replicate means from 3 independent experiments in B-D, F-H.

A

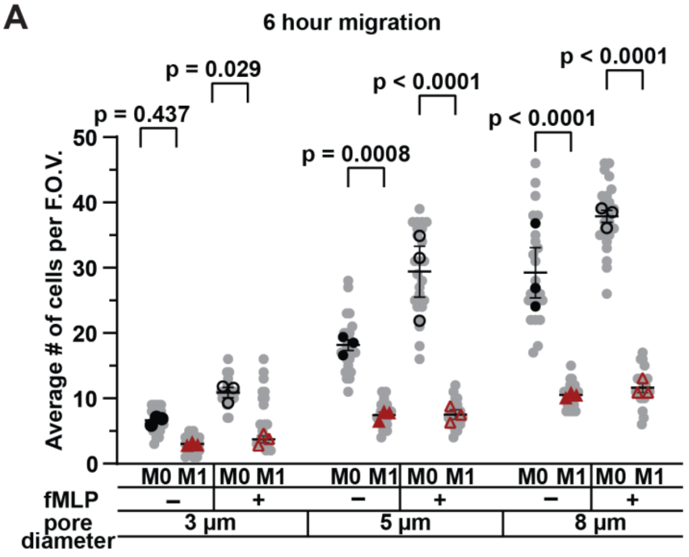


B

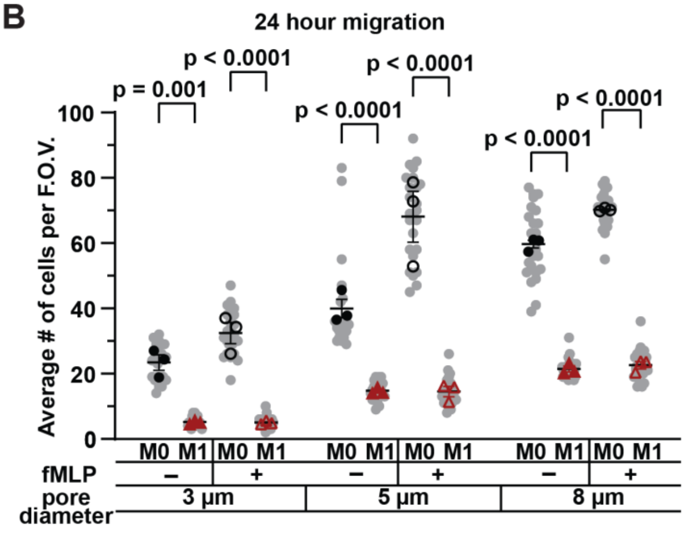


C

| 6hours | | | | | |
|---|---|---|---|---|---|
| 3 µm | P value | 5 µm | P value | 8 µm | P value |
| M0 vs. M0 + fMLP | 0.3004 | M0 vs. M0 + fMLP | 0.0004 | M0 vs. M0 + fMLP | 0.0068 |
| M0 vs. M1 | 0.4371 | M0 vs. M1 | 0.0008 | M0 vs. M1 | <0.0001 |
| M0 vs. M1 + fMLP | 0.6208 | M0 vs. M1 + fMLP | 0.0009 | M0 vs. M1 + fMLP | <0.0001 |
| M0 + fMLP vs. M1 | 0.0141 | M0 + fMLP vs. M1 | <0.0001 | M0 + fMLP vs. M1 | <0.0001 |
| M0 + fMLP vs. M1 + fMLP | 0.0286 | M0 + fMLP vs. M1 + fMLP | <0.0001 | M0 + fMLP vs. M1 + fM | <0.0001 |
| M1 vs. M1 + fMLP | 0.9896 | M1 vs. M1 + fMLP | >0.9999 | M1 vs. M1 + fMLP | 0.9635 |

D

| 24 hours | | | | | |
|---|---|---|---|---|---|
| 3 µm | P value | 5 µm | P value | 8 µm | P value |
| M0 vs. M0 + fMLP | 0.1234 | M0 vs. M0 + fMLP | <0.0001 | M0 vs. M0 + fMLP | 0.0582 |
| M0 vs. M1 | 0.0005 | M0 vs. M1 | <0.0001 | M0 vs. M1 | <0.0001 |
| M0 vs. M1 + fMLP | 0.0005 | M0 vs. M1 + fMLP | <0.0001 | M0 vs. M1 + fMLP | <0.0001 |
| M0 + fMLP vs. M1 | <0.0001 | M0 + fMLP vs. M1 | <0.0001 | M0 + fMLP vs. M1 | <0.0001 |
| M0 + fMLP vs. M1 + fMLP | <0.0001 | M0 + fMLP vs. M1 + fMLP | <0.0001 | M0 + fMLP vs. M1 + fML | <0.0001 |
| M1 vs. M1 + fMLP | >0.9999 | M1 vs. M1 + fMLP | >0.9999 | M1 vs. M1 + fMLP | 0.9913 |

**Supplementary Figure 7. Addition of chemoattractant does not increase M1 migration though confined or un-confined pores. (A**) Quantification of the number of cells that completed migration through transwell membranes by 6 hours with the addition of fMLP or vehicle control. (**B**) Quantification of the number of cells that completed migration through transwell membranes by 24 hours with the addition of 100 nM fMLP or vehicle control. **(C, D)** Table showing statistics of (**C**) 6-hour and (**D**) 24-hour migration using a two-way ANOVA performed on the replicate means with Tukey's multiple comparison test. Quantification is represented as horizontal lines at the mean ± SEM, with replicate means plotted in large black circles (M0) or red triangles (M1), and individual measurements shown as small grey circles.
